# Convergent molecular signatures of ageing and injury in the peripheral nervous system

**DOI:** 10.64898/2026.02.09.704884

**Authors:** Dario Lucas Helbing, Michael Reuter, Emilio Cirri, Therese Thuy Dung Dau, Joanna M. Kirkpatrick, Amy Stockdale, Alexander Schulz, Nova Oraha, Leopold Bohm, Philipp Koch, Leonie Karoline Stabenow, Alexander Weuthen, Nadja Gebert, Martin Walter, Alessandro Ori, Karl Lenhard Rudolph, Reinhard Bauer, Helen Morrison

## Abstract

Peripheral nervous system (PNS) ageing is marked by structural and functional decline, yet it remains unclear whether ageing constitutes a distinct biological programme or reflects a chronic injury-like state. To address this, we performed an unbiased, comparative molecular analysis of PNS ageing, neuroprotective dietary restriction (DR), and nerve injury. We conducted transcriptomic and proteomic profiling of peripheral nerves from young, old and geriatric mice fed ad libitum or subjected to long-term DR, and proteomics of nerves collected at multiple time points following injury. Age-associated molecular changes followed both linear and non-linear trajectories, and DR partially attenuated these ageing-related alterations. Notably, ageing-and injury-induced proteomic signatures showed considerable similarities, supporting the concept that an aged nerve resembles an injured nerve. Together, our study provides the most comprehensive molecular resource of PNS changes during ageing, DR, and injury, enabling the definition of key molecular signatures underlying PNS physiology. All datasets are integrated into the “PNS-omics Viewer”, a Shiny web application designed to facilitate data mining of the herein presented datasets (*tba*).

## Main

Peripheral nervous system (PNS) ageing leads to structural and functional changes that influence the regenerative potential and maintenance of the PNS. This results – in addition to general age-related decline in nerve function, such as reduced vibration sense, electrophysiological alterations, and absent tendon reflexes^1^, – in age-associated peripheral neuropathies with no other identifiable cause, i.e. idiopathic neuropathies, that may account for up to 39% of all cases in patients aged ≥80 years^2^. However, the identification of underlying age-associated peripheral nerve-intrinsic changes to determine pathways and biological functions that could be targeted to restore tissue homeostasis, maintenance and regenerative capacity of old nerves, has been inconclusive thus far. Therefore, using a hypothesis-free approach, we aimed to understand how peripheral nerves change during ageing, and used RNA-sequencing and mass-spectrometry to characterise molecular changes in old (20 months) and geriatric (24 months) nerves as compared to young (3 months) nerves (**Figure 1A**).

**Fig. 1.**
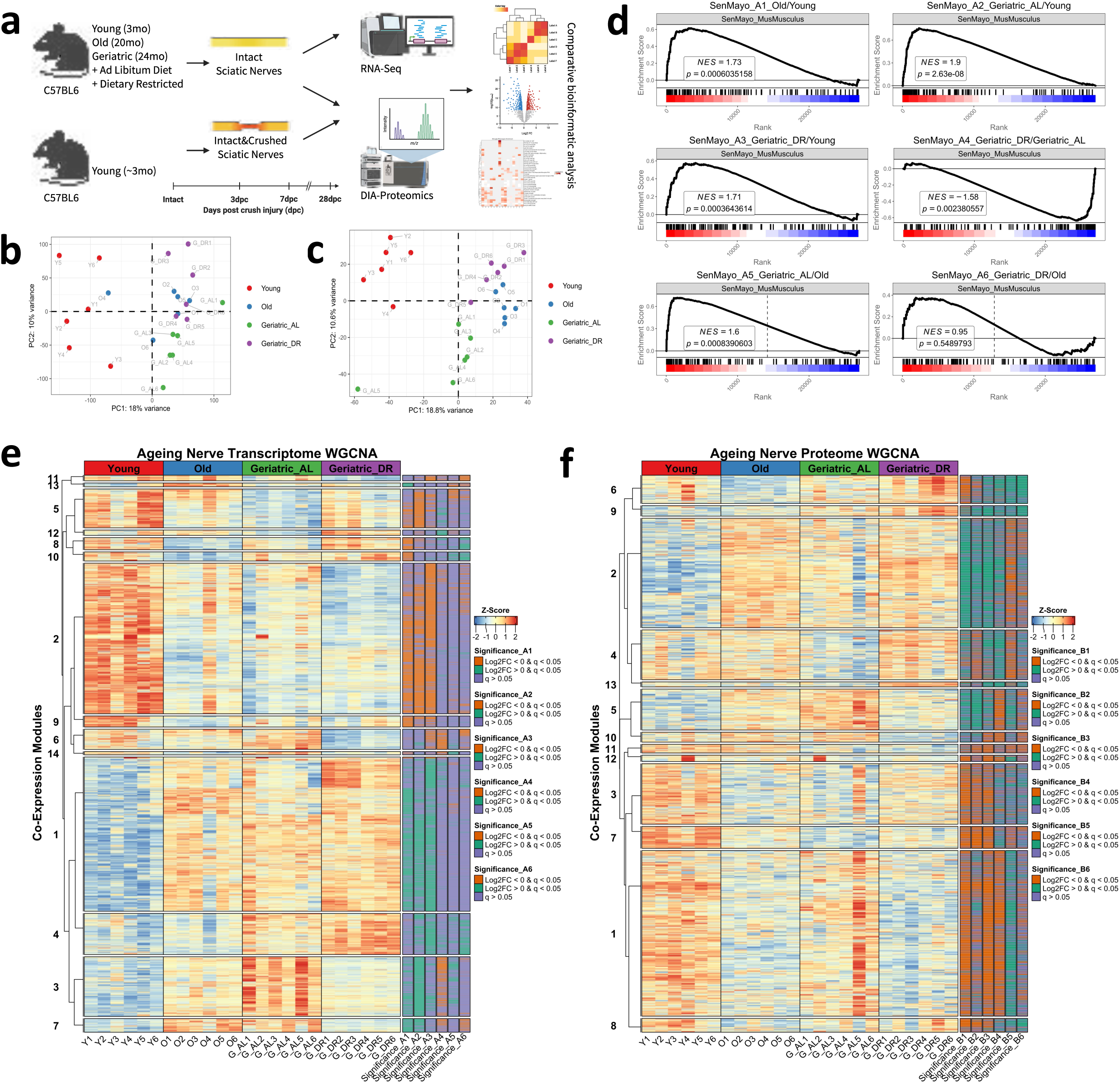
I Transcriptomic and proteomic analysis of peripheral nerve ageing and long-term dietary restriction. **A** Schematic experimental workflow for integrative analysis of peripheral nerve transcriptomic and proteomic changes during ageing and long-term dietary restriction (LT-DR) (n = 6 per group), as well as proteomic analysis of peripheral nerves during nerve degeneration and regeneration after experimental crush injury (n = 4 – 11 per post-injury group). **B, C** Principal component analysis (PCA) of PNS ageing & LT-DR transcriptome (**B**) and proteome (**C**), the different experimental groups (n = 6 per group) are coloured as depicted in the legend on the right side of the PCA plots. **D** Gene set enrichment analyses in the ageing & LT-DR transcriptome of the SenMayo senescence geneset^25^ for all different comparisons among all experimental groups 1 = O/Y, 2 = G_AL/Y, 3 = G_DR/Y, 4 = G_DR/G_AL, 5 = G_AL/O, 6 = G_DR/G_AL. **E, F** Identification of different co-expression modules that represent genes/transcripts (**E**) and proteins (**F**) that are co-regulated across different clusters with weighted gene co-expression network analysis (WGCNA). Either transcript-or protein-wise z-score normalized intensities are shown and on the right side an annotation of Log2 FC and q-value levels are depicted (Log2 FC≥ 0 in green or Log2 FC≤ 0 in orange, indicating significant up-or downregulation in the comparisons A1/B1-A6/B6 (q-value < 0.05) respectively or a q-value > 0.05 in purple). Abbreviations A1-A6/B1-B6 represent the different comparisons among all experimental groups, 1 = O/Y, 2 = G_AL/Y, 3 = G_DR/Y, 4 = G_DR/G_AL, 5 = G_AL/O, 6 = G_DR/G_AL.

PNS ageing is characterised by axonal loss, myelin abnormalities and an overall shift of the cellular composition and microenvironment^3–5^. In particular, the glia cells of the PNS, Schwann cells, display differences in their activity and differentiation state^4,5^ that may also account for their impaired phagocytic activity in old age^4,5^. Furthermore, “Inflammageing”, defined by an increased, low-grade presence of inflammation and especially innate immune cells, i.e. macrophages, in peripheral tissues such as old, intact nerves as well as the impaired capacity to fulfill core macrophage functions (i.e. phagocytosis of myelin debris), and an altered response to any noxae contributes to age-related peripheral nerve pathology and limited regenerative capacity^6–9^. Markedly, the increased inflammation in old, intact nerves – that reminds of the inflammation seen in young nerves after a peripheral nerve injury – and a subsequently amplified immune response to e.g. PNS injury appears to be a major contributing factor to the impaired regenerative capacity of the old PNS ^7,9^. Moreover, in old intact nerves, Schwann cells can be observed in an injury-like, non-functional repair mode, and it remains unclear if Inflammageing contributes to this response^9^. Interestingly, Yuan et al. demonstrated that 24-month-old mice exhibit pronounced peripheral nerve degeneration (which begins around 18 months of age, with no such alterations observed in 12-month-old mice) and increased macrophage accumulation after 18 months. Targeting macrophages by depleting them before 18 months led to improved nerve structure at 24 months, indicating that major age-related changes may start to occur around 18-20 months^7^. These observations are supported by Comfort et al. who performed transcriptomic analysis only and detected first transcriptional changes in sciatic nerves at 18 months^10^, as well as Walsh et al., who measured nerve conduction velocity (NCV) in mice aged 4–32 months and only observed a decline of NCV in mice starting at 20 months of age^11^, and Krishnan et al., who noted signs of impaired proteostasis and axonal transport from about 18 months onward^12^. However, the major driving factors that define and determine the inflammatory state, as well as overall molecular alterations of the PNS during ageing, remain largely unknown.

In general, dietary restriction (DR) and, more specifically, caloric restriction (CR) are considered to be among the most effective interventions for extending the lifespan and healthspan of various organisms, including mice and humans, by preserving homeostasis and slowing down the ageing of multiple organs^13^. Although the importance of the PNS in regulating organ homeostasis and ageing across several tissues has become more widely recognised in recent years^14–21^, the effects of DR/CR on PNS ageing itself have not yet been studied systematically. Nevertheless, previous functional studies have identified that caloric restriction attenuates age-related structural and functional decline in the peripheral nervous system (PNS), particularly in geriatric mice (24+ months). One study by Walsh et al. could demonstrate that life-long CR preserved sciatic NCV in geriatric mice^11^, and in another study, life-long CR attenuated motor neuron loss in geriatric (24 months) mice, which otherwise started at around 18 months and was severe by 24 months in mice fed an *ad libitum* diet^22^. In addition, Rangaraju et al. noted that life-long CR preserved myelin degeneration (which was otherwise not detectable at 18 months yet) and Schwann cell phenotype in geriatric mice^23^. Taken together, caloric restriction consistently preserves peripheral nerve health and function in old and geriatric mice, with the greatest effects seen with lifelong or early-onset intervention – however, the molecular effects of CR on the PNS have not been investigated yet in a systematic manner.

In this study, we extend previous reports and investigate with an unbiased, comparative approach how ageing shapes peripheral nerve transcriptome and proteome trajectories from young to old and geriatric age. Furthermore, we aimed to identify the main biological functions and pathways influenced by long-term dietary/caloric restriction, as a known neuroprotective intervention for healthy PNS ageing. Lastly, we also wanted to examine peripheral nerve injury profiles during PNS de– and regeneration and compare age– and injury-induced proteomic shifts (**Figure 1A**). Our analyses reveal a stepwise molecular PNS ageing process that is markedly slowed by DR. We identify convergent metabolic and inflammatory networks – regulated at both transcriptional and translational levels – that underlie peripheral nerve ageing and its attenuation by DR. Remarkably, we identified parallels between metabolic and inflammatory remodelling during ageing and PNS injury, which leads us to the conclusion that, at a molecular level, an old nerve is an injured nerve. We believe that this study represents the most comprehensive molecular resource on peripheral nerve ageing, injury, and dietary modulation to date, and serves as a valuable resource for understanding the molecular physiology of the ageing peripheral nervous system, and to identify new strategies to preserve peripheral nerve function and regenerative potential at old age.

## Extended data figure legends

**Fig. ED1.**
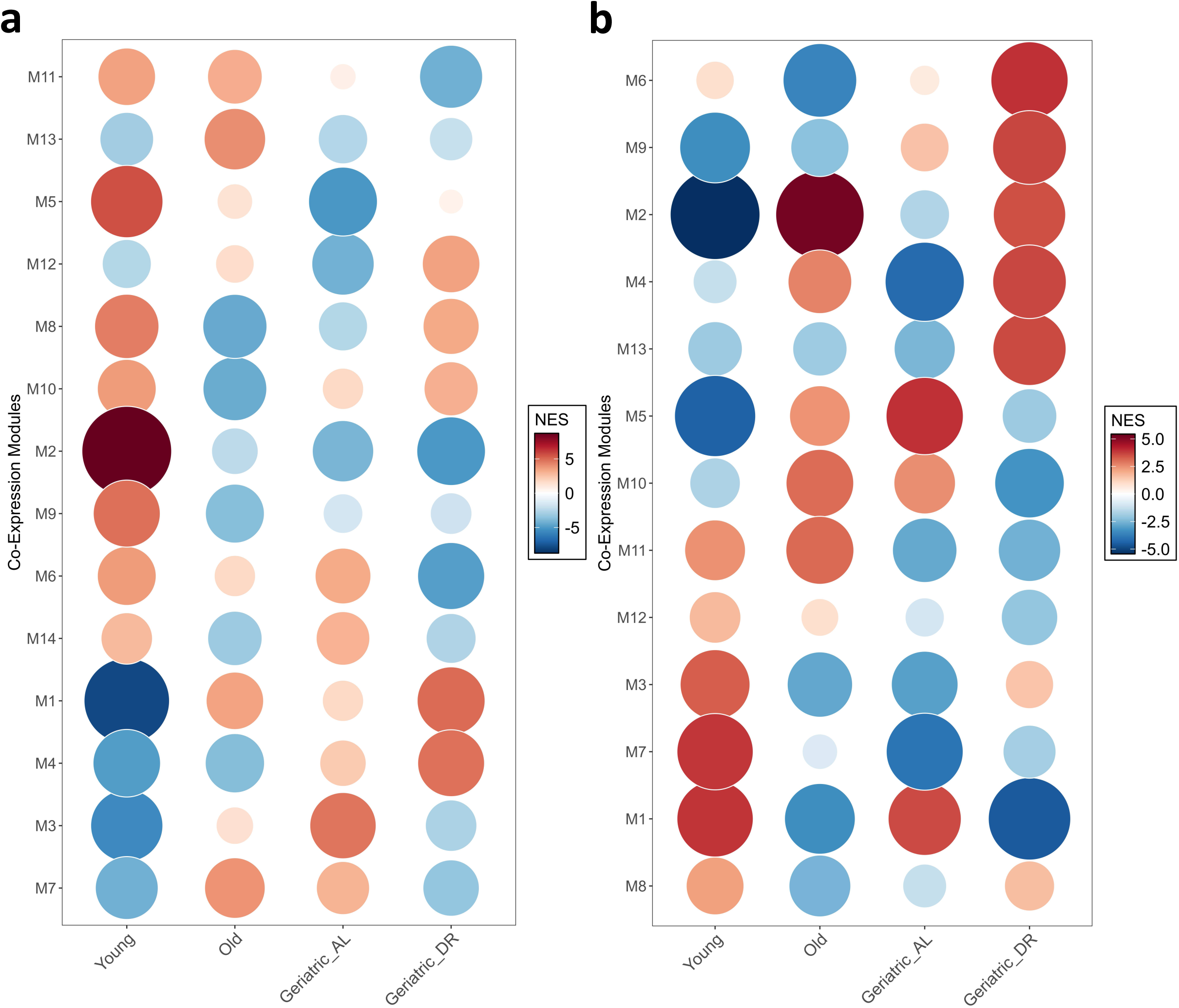
I Enrichment of co-expression modules across experimental cohorts. **A, B** Numerical enrichment scores for the identified co-expression modules shown in figure 1 across all different experimental groups in the transcriptome (**A**) or proteome (**B**) visualizes patterns of ageing trajectories and their regulation by LT-DR.

**Fig. ED2.**
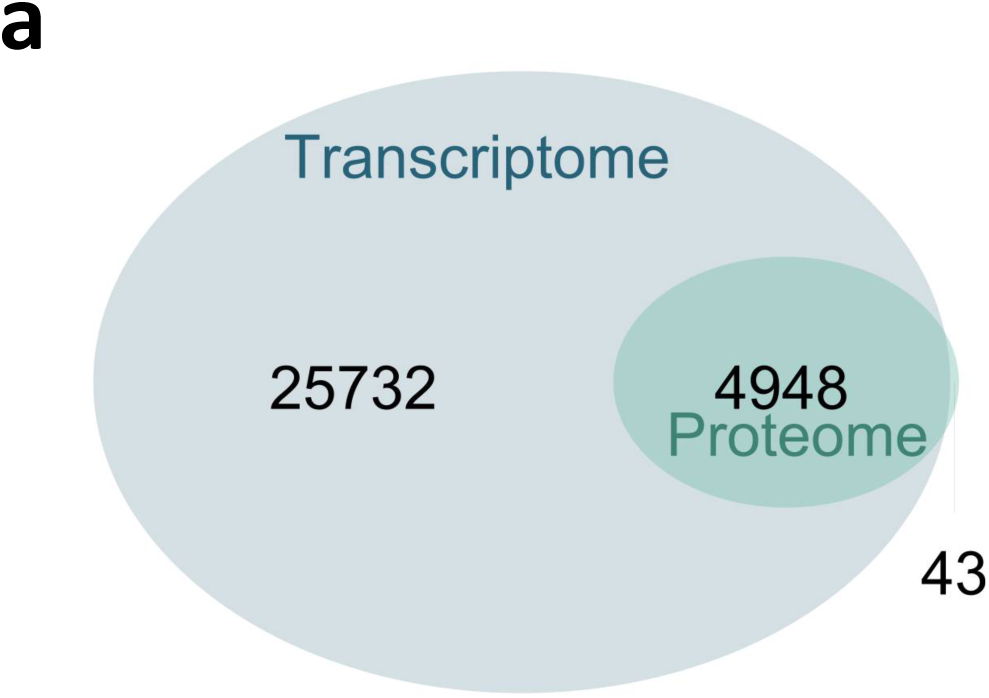
I Transcriptome-proteome overlap. **A** Venn diagram showing the number of transcripts detected only in the transcriptome (n = 25732), the number of gene products for which both the transcript and the protein were detected (n = 4948), and the number of proteins detected only in the proteome (n = 43).

**Fig. ED3.**
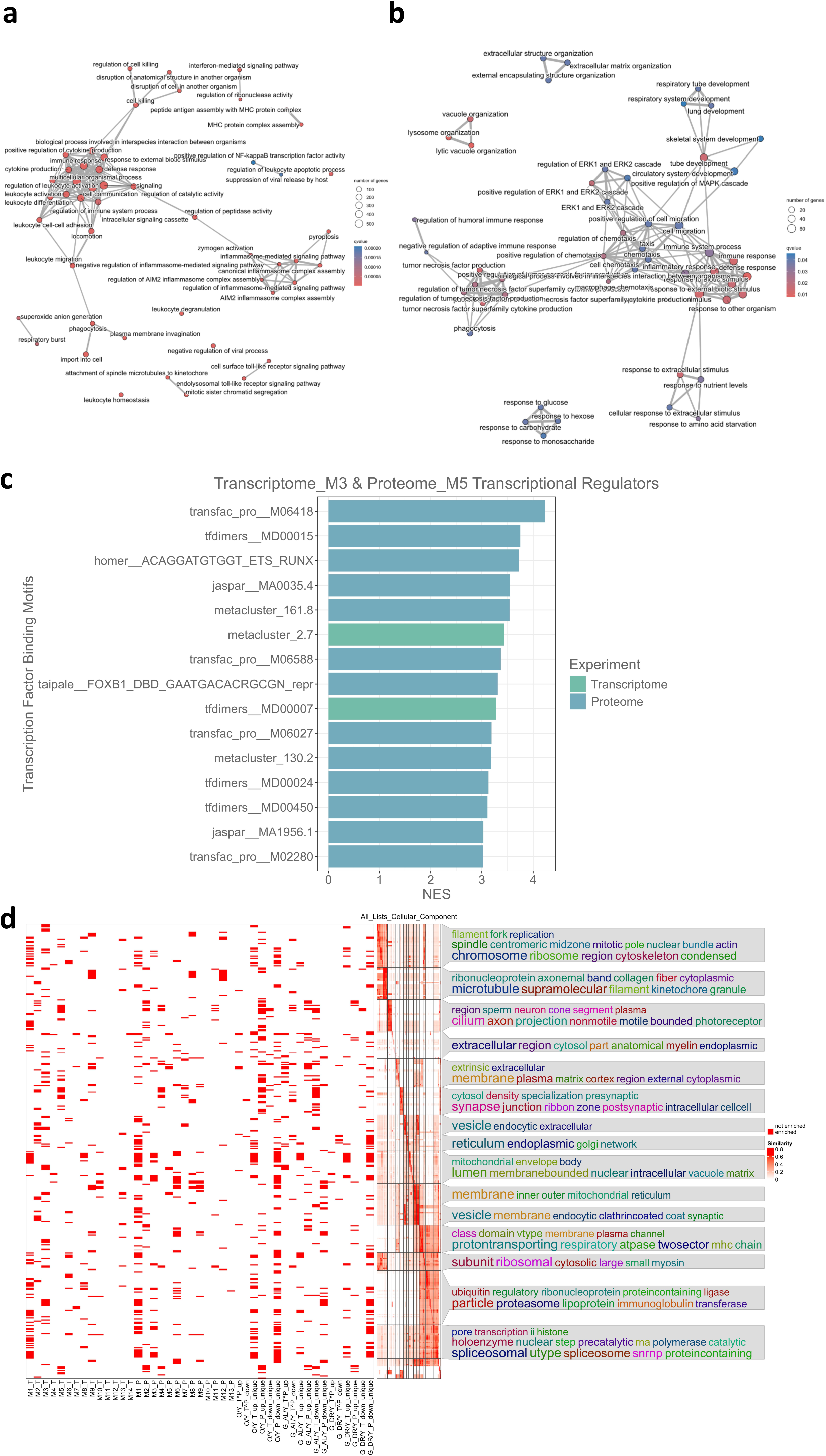
I Inflammatory modules and cellular component changes during peripheral nerve ageing and long-term dietary restriction. **A, B** Enrichment map as network visualization of biological function gene ontology terms enriched in the M3 cluster in transcriptome (**A**) and M5 cluster in the proteome (**B**). Gene ontology terms with larger overlap of annotated genes/proteins do cluster together and the number of annotated genes to the respective gene ontology term as well as the q-value of the over-representation analysis for the respective GO-term are shown on a colour scale. **C** Numerical enrichment scores of the top 15 enriched transcription factor binding motifs in either the M3 cluster of the transcriptome or the M5 cluster of the proteome are shown. A full list of identified motifs in M3_T and M5_P in addition to the predicted transcriptional regulators is shown **supplementary table 1** and a full list of all identified motifs and transcriptional regulators is provided in **supplementary dataset 2**. **D** Gene ontology enrichment analysis of cellular component GO-terms for all WGCNA modules and significantly up-/downregulated genes/proteins in the individual experimental comparisons 1-6 (1 = O/Y, 2 = G_AL/Y, 3 = G_DR/Y, 4 = G_DR/G_AL, 5 = G_AL/O, 6 = G_DR/G_AL) using semantic similarity analysis, methodically similar to Fig. 2

**Fig. SED.**
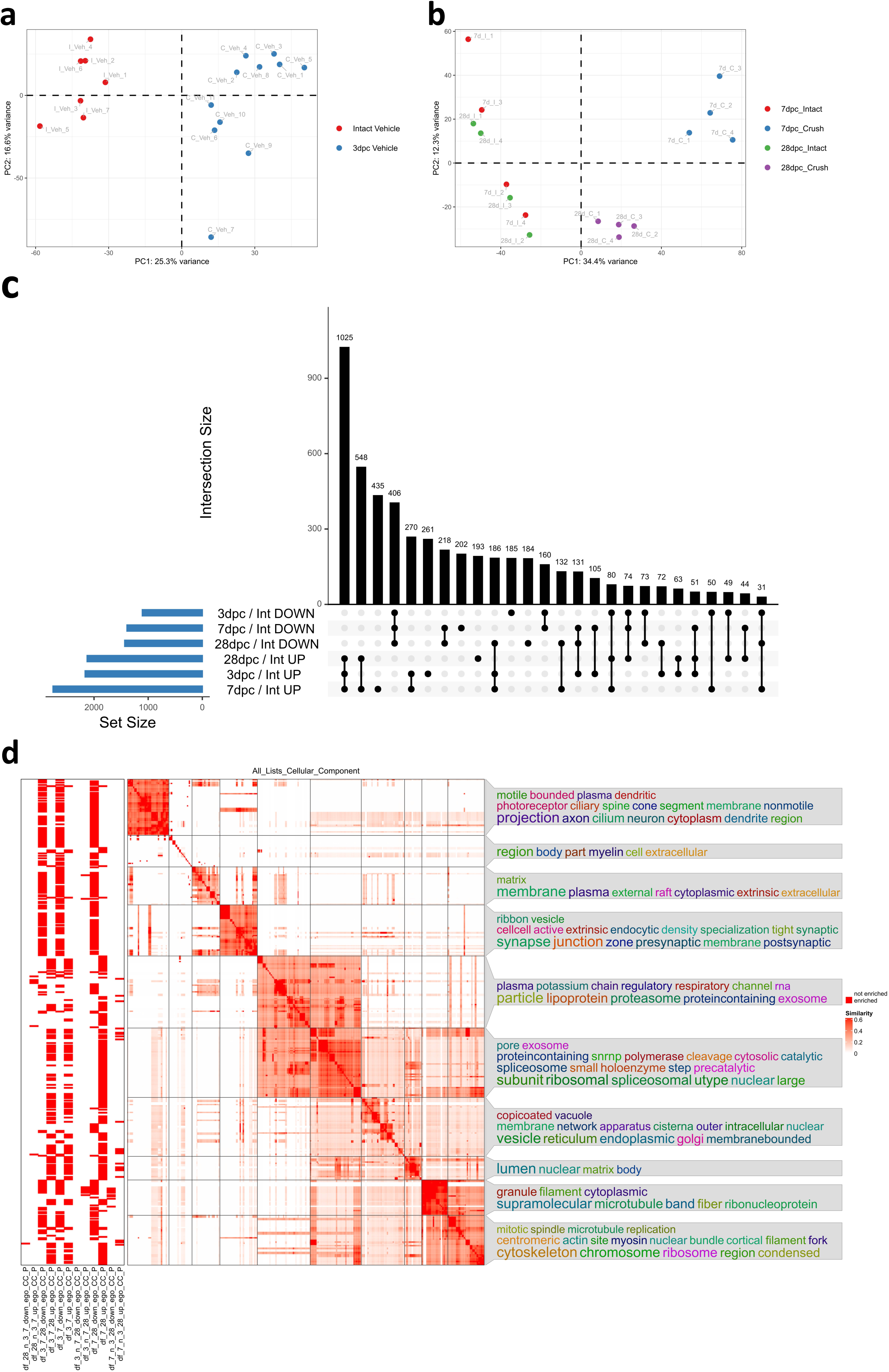
I Functional analysis of PNS crush injury proteomics datasets. **A, B** Principal component analysis (PCA) of 3dpc nerve injury proteome (**A**) and 7&28dpc nerve injury proteome (**B**), the different experimental groups (n = 4-11 per group) are coloured as depicted in the legend on the right side of the PCA plots. **C** Upset plots showing the number of shared differentially regulated proteins (depicted as intersection size) between the different significantly up– and downregulated proteins 3, 7 and 28dpc. The size of the respective module or gene/protein list is displayed on the right side of the plot (labelled as set list). **D** Gene ontology enrichment analysis of cellular component GO-terms for all individual time-specific injury response proteome profiles using semantic similarity analysis, methodically similar to Fig. 2. Column names for the different protein sets are described in detail in the methods.

**Fig. ED5.**
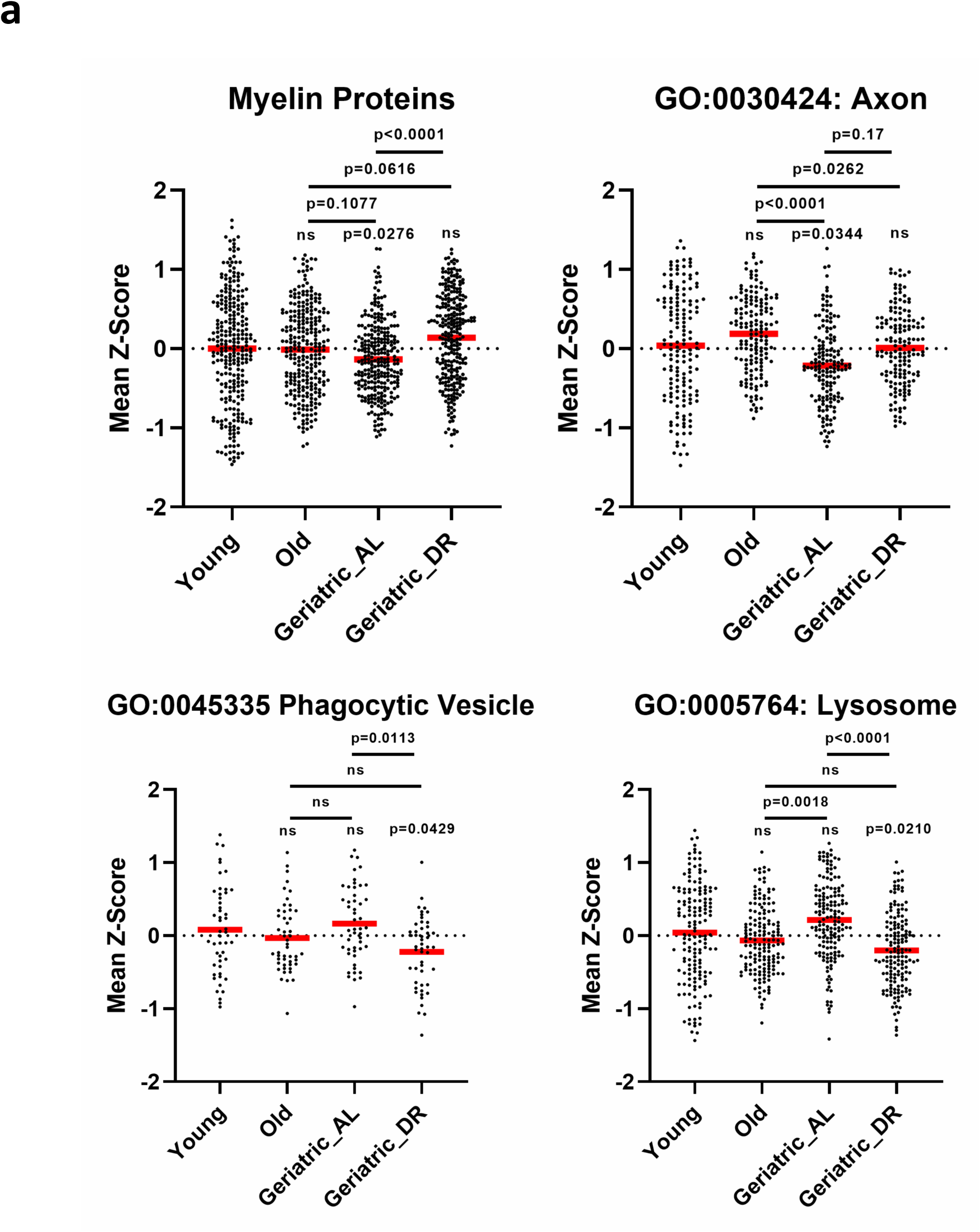
I Analysis of myelin, axonal and phagocytic/lysosomal vesicle proteins during PNS ageing and LT-DR. **A** Mean z-scores (across n=6 biological replicates) of either all myelin proteins as reported in Helbing et al., 2023^45^ and all axonal, phagocytic vesicle and lysosomal proteins that are annotated to the respective GO-terms and were detected in this dataset. Friedman tests for myelin, lysosomal and axonal proteins and a repeated-measures one-way ANOVA for phagocytic vesicle proteins were performed, each with p < 0.05– For Friedman tests, post-hoc Dunn’s multiple comparisons tests were performed and for the RM-ANOVA post-hoc paired t-tests were performed and p-values were FDR-corrected (Benjamini-Hochberg method). Shown are the mean (red lines) ± SEM.

## Supplementary tables

**Supplementary Table 1:**
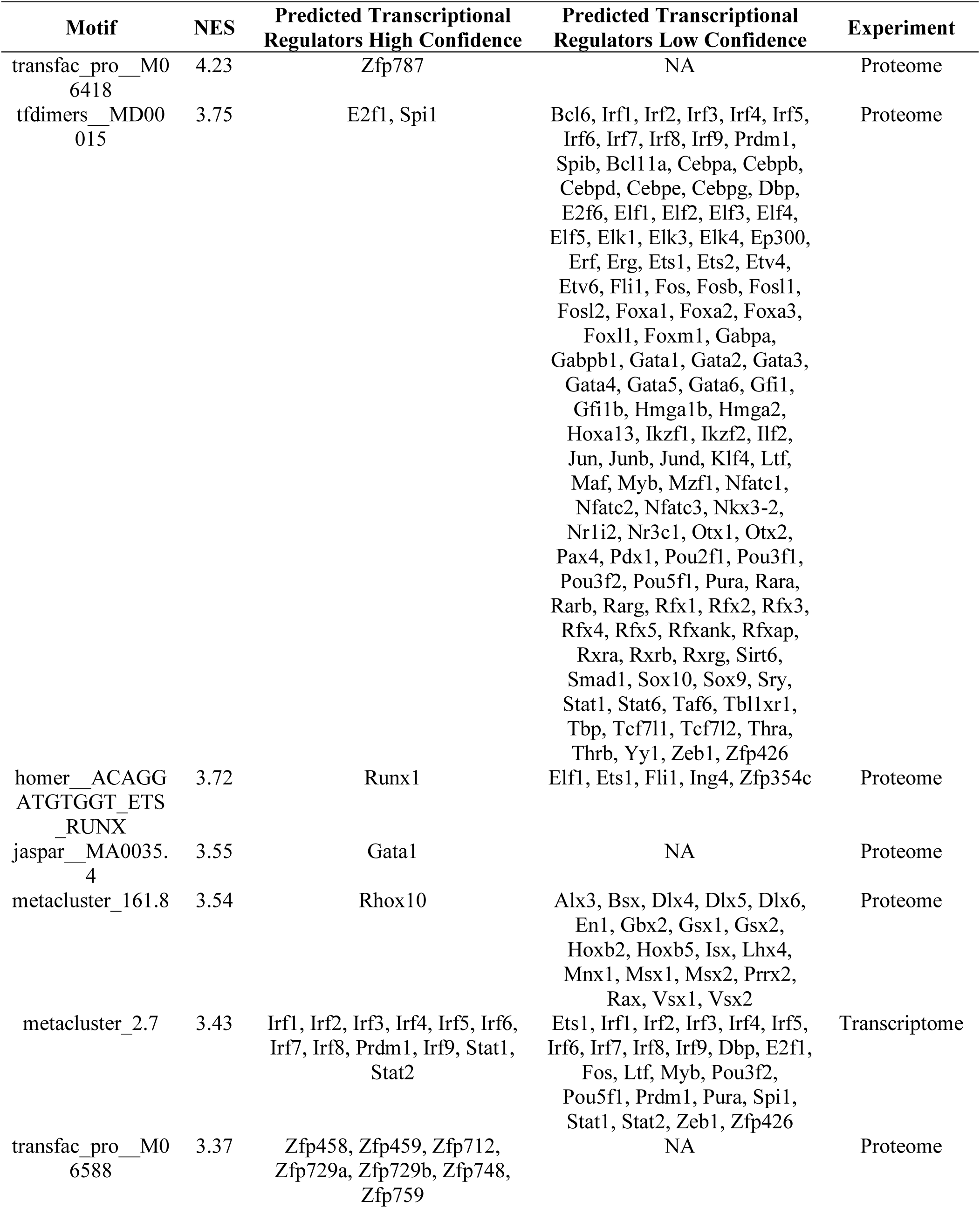
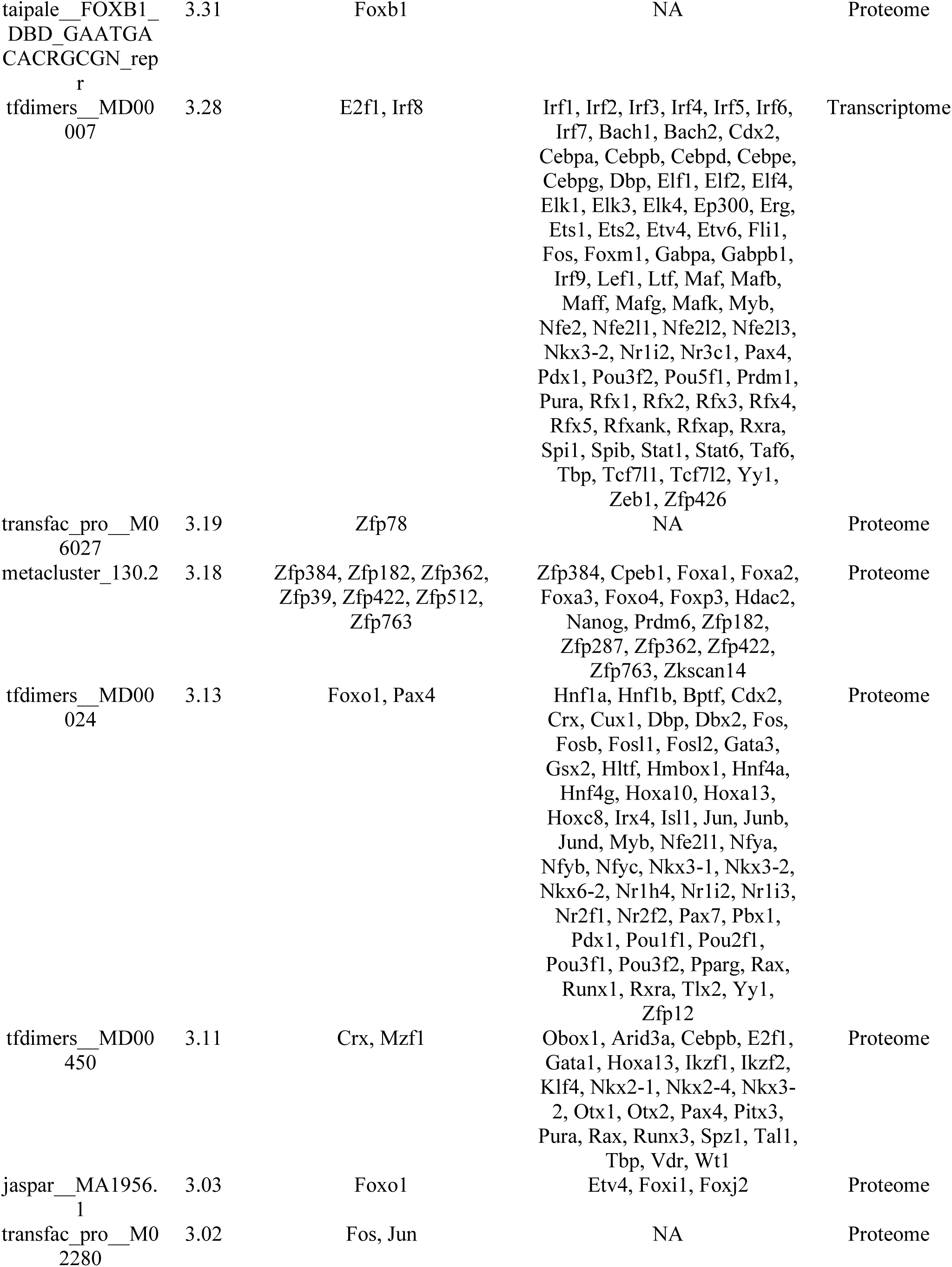
Enriched Transcription Factor Binding Motifs and Predicted Transcriptional Regulators in the Modules M3 (Transcriptome) and M5 (Proteome)

**Supplementary Table 2:** Spearman’s ρ of correlation pairs proteomics datasets.

| Dataset | G_AL/O | C/I_7dpc | C/I_3dpc | C/I_28dpc | G_DR/O | G_AL/Y | G_DR/G_AL | O/Y | G_DR/Y |
| --- | --- | --- | --- | --- | --- | --- | --- | --- | --- |
| G_AL/O | NA | 0.376725783 | 0.392191593 | 0.390203449 | 0.305372289 | 0.198397441 | -0.557559627 | -0.526762308 | -0.291469118 |
| C/I_7dpc | 0.376725783 | NA | 0.57792598 | 0.618208624 | -0.013025278 | 0.078402561 | -0.37114882 | -0.206903304 | -0.241177946 |
| C/I_3dpc | 0.392191593 | 0.57792598 | NA | 0.400717608 | -0.03981932 | 0.039024871 | -0.398506714 | -0.248946971 | -0.291976444 |
| C/I_28dpc | 0.390203449 | 0.618208624 | 0.400717608 | NA | 0.062792453 | 0.007038136 | -0.300665348 | -0.259618185 | -0.230225397 |
| G_DR/O | 0.305372289 | -0.013025278 | -0.03981932 | 0.062792453 | NA | -0.154339 | 0.516176942 | -0.366389314 | 0.28992574 |
| G_AL/Y | 0.198397441 | 0.078402561 | 0.039024871 | 0.007038136 | -0.154339 | NA | -0.30393137 | 0.641826254 | 0.557010418 |
| G_DR/G_AL | -0.557559627 | -0.37114882 | -0.398506714 | -0.300665348 | 0.516176942 | -0.30393137 | NA | 0.142267556 | 0.520007205 |
| O/Y | -0.526762308 | -0.206903304 | -0.248946971 | -0.259618185 | -0.366389314 | 0.641826254 | 0.142267556 | NA | 0.706701545 |
| G_DR/Y | -0.291469118 | -0.241177946 | -0.291976444 | -0.230225397 | 0.28992574 | 0.557010418 | 0.520007205 | 0.706701545 | NA |

## Results

### Comparative and functional analysis of transcriptome and proteome changes during PNS ageing and dietary restriction

In order to comprehensively assess peripheral nervous system changes with ageing and dietary restriction as neuroprotective intervention, we isolated RNA and proteins from the same sciatic nerves from young (3 months, “Y”), old (20 months, “O”) and geriatric mice (24 months) fed either an *ad libitum* diet (“G_AL”) or that were kept long-term dietary/caloric restricted from 4 to 24 months (“G_DR”) and performed transcriptomic and proteomic profiling, using RNA-sequencing and mass-spectrometry based proteomics, respectively. In addition, we performed proteomics on sciatic nerves at 3 days post nerve crush injury (3dpc), 7dpc and 28dpc and compared time-point specific injury profiles with ageing profiles to characterize similar and distinct features between ageing– and injury responses in the PNS (**Figure 1A**). To categorize global transcriptomic and proteomic changes in the PNS throughout ageing and DR, we performed a principal component analysis on the total datasets. In both datasets, a clear separation along the first principal component is observed between young nerves and all old/geriatric nerves, indicating a strong impact of ageing on both, the transcriptome and proteome of the PNS (**Figure 1B, C**). Furthermore, while evident in both datasets, particularly in the proteome (**Figure 1C**), an age-effect is also discernible on the second principal component, with O and G_AL nerves distinctly separated from young nerves in a step-wise manner and G_DR nerves partially “rescued”, i.e. on the same scale across PC2 as young nerves (**Figure 1B, C**). To evaluate induction of a senescence phenotype as an hallmark ageing phenotype^24^, we used the SenMayo geneset, which has been validated to identify senescent cells on transcriptome level across tissues and species^25^ to perform a gene set enrichment analysis (GSEA) on our transcriptomic dataset. The SenMayo geneset was markedly enriched in old and even stronger in G_AL nerves, indicating a stepwise induction of senescence transcriptomic programs during PNS ageing (**Figure 1D**). It is interesting to note that, although G_DR nerves exhibited an upregulation of SenMayo genes when compared to young nerves, in comparison to G_AL nerves, the SenMayo genes were downregulated, with no detectable difference as compared to old nerves. This indicates, based on the analysis of senescence marker genes, that there is slowed ageing on transcriptome level in G_DR nerves (**Figure 1D**). As expected and initial hierarchical clustering analyses indicated (data not shown), ageing trajectories and modification of these by dietary restriction show only partially linear dynamics and follow mostly a non-linear manner, which has been shown for most organs before in human and mice^26^. Therefore, we employed weighted gene co-expression/correlation network analysis to identify modules/clusters of highly correlated transcripts/proteins (**Figure 1E, F**). In sum, 14 clusters could be identified in the ageing nerve transcriptome dataset and 13 clusters were identified in the ageing nerve proteome dataset, with heterogeneous dynamics that are visible in the heatmaps (**Figure 1E, F**) as well as the plots depicting the numerical enrichment scores of the individual modules across all experimental cohorts (**Extended Data Figure 1A, B**). Several clusters showed a linear, stepwise up-/downregulation (e.g. M3_transcriptome_, M5_proteome_ (upregulation); M5_transcriptome_, M7_proteome_) with “rescue” in the G_DR group, i.e. similar transcript/protein profiles as compared to old nerves. However, multiple other clusters can be observed, showing patterns of up-/downregulation in old and G_DR nerves, but similar levels to young nerves in G_AL nerves (e.g. M1_proteome_, M2_proteome_), or exhibiting an increase/decrease during the process of ageing, with higher transcript/protein abundance in G_DR nerves (e.g. M9_proteome_, M6_proteome_, M10_transcriptome_, M4_transcriptome_). This indicates that dietary restriction exerts an effect on the physiological ageing process that partially differs from a simple “slowdown” of ageing processes (**Figure 1E, F, Extended Data Figure 1A, B**). Although most of the observed proteomic patterns resemble the transcriptome patterns and vice versa, different profiles were also visible, which is why we compared the overlap of all the gene product modules detected in both experiments (4948 transcripts/proteins, **Extended Data Figure 2A**).

It is notable that only some of the largest overlaps were between modules that displayed similar patterns (e.g. M1_proteome_/_M2transcriptome_) while several overlaps occurred between modules that displayed opposite patterns. (e.g. M1_proteome_/M1_transcriptome_, M2_proteome_/M2_transcriptome_, M3_proteome_/M2_transcriptome_). However, amongst the smaller overlaps were multiple modules with similar patterns between transcriptome and proteome profiles (e.g. M5_proteome_/M_3transcriptome_ and M4_transcriptome_/M6_proteome_) (**Figure 2A**). To determine the changes induced by ageing and DR in the PNS, we conducted gene ontology (GO) enrichment analyses for all transcriptome and proteome modules and comparisons, focusing on biological processes (BP) and cellular components (CC), in addition to KEGG pathway enrichment analyses. Similar enrichment profiles are especially visible for M3_transcriptome_, M1_proteome_, M5_proteome_ and the list of upregulated transcripts and proteins in G_AL/Y as well as partially upregulated proteins in O/Y, with multiple immune– and inflammation-related GO-terms being enriched. Specifically, “interferon”, “interferonalpha”,”interferonbeta” and also “nfkappab” were keywords that appeared several times amongst the in the aforementioned lists enriched GO_BP terms, reflecting the stepwise induction of inflammatory processes during ageing in the PNS and it’s apparent prevention by LT-CR (**Figure 2B** and **Supplementary Dataset 3**). To explore these modules in more detail, we plotted an enrichment map of the enriched biological process GO-terms (full names). Indeed, nearly only GO-terms related to immune activation, cytokine production, inflammasome but also phagocytosis and specifically, ERK, NF-κB and interferon-mediated signalling were overrepresented in these modules (**Extended Figure 3A, B**). To further substantiate these results, we analyzed transcription factor binding motifs in all the transcript and protein lists of all the modules, focusing on the M3_transcriptome_ and M5_proteome_ modules (**Extended Figure 3C**, **Supplementary Table 1, Supplementary Dataset 2**). Amongst the most prominent transcriptional regulators were e.g. Zfp787, a transcriptional regulator protein with so far unknown PNS functions and multiple transcriptional regulator proteins with predominant expression in immune cells (e.g. Spi1, Runx1, Gata1) but also regulators with known functions in the PNS, especially Schwann cells (e.g. Foxo1, Fos, Jun). Moreover, when all regulators are considered, the largest number of predicted transcriptional regulators functionally belong to the interferon response or regulatory system (e.g. the IRF and STAT families), which, similar to the GO enrichment results, may indicate an interferon-response immune state in the ageing PNS (**Extended Figure 3C**, **Supplementary Table 1**). In M9_transcriptome_ and M12_proteome_, modules that contained primarily with ageing downregulated transcripts and proteins, were a large number of biological processes enriched that are associated to “development”, “differentiation”, “morphogenesis”, “angiogenesis”, “transport” GO-terms, which likely reflects ongoing developmental processes in the young PNS (**Figure 2B)**. However, in the same modules, in addition to M3_transcriptome_, “vesicle”, “phagocytosis” GO-terms were enriched, indicating potential differences in clearance processes (**Figure 2B)**. A large number of biological process GO-terms related to “splicing”, “transcription”, “mrna” and metabolism related to splicing and translation (**Figure 2B)** and the GO_CC terms “spliceosomeal”, “polymerase” and “ribosomal” (**Extended Figure 3D**) were enriched predominantly in the clusters that show opposite patterns between transcriptome and proteome, M1_transcriptome_ and M1_proteome_ (**Figure 1E**, **F**). Moreover, two GO-term clusters related to primarily glucose metabolism (“metabolic”, “-phosphate”, “monocarboxylic”) and lipid metabolism showed enrichment amongst several transcript/protein lists, specifically M6,9,10_proteome_ and M9,M10,M4_transcriptome_, as well as in the lists of by dietary restriction upregulated transcripts/proteins and uniquely by DR upregulated proteins (G_DR/Y), indicating profound effects of lifelong caloric restriction on glucose and lipid metabolism in the PNS (**Figure 2B**). Concomitantly, cellular component GO-terms containing mitochondrial proteins, also specifically belonging to the inner/outer membrane, were enriched in the same lists (**Extended Figure 3D**). In order to further investigate changes in metabolism, signal transduction and immune activation, we also performed an enrichment analysis for KEGG pathways. We observed primarily a cluster of 3 modules with upregulation in G_DR nerves – M6,M9_proteome_ and M10_transcriptome_ – to show similar enrichment patterns in metabolic pathways related to glucose catabolism including glycolysis and oxidative phosphorylation, while the latter was also enriched in the uniquely downregulated proteins in G_AL and O groups (**Figure 2C**). Furthermore, lipid metabolism, both fatty acid biosynthesis and degradation were enriched in the same 3 modules and steroid biosynthesis was enriched in the lists of downregulated transcripts and proteins in all ageing cohorts (O, G_AL and G_DR). Moreover, the KEGG pathway “Valine, leucine and isoleucine degradation” was enriched in M6,M9_proteome_ and M10_transcriptome_ as well as the in G_DR nerves upregulated transcripts and proteins. It is worth noting that well-known ageing-associated KEGG pathways “taurine and hypotaurine metabolism”^27^, “Glutathione metabolism”^28^ and additionally “beta-Alanine metabolism”^29^ were enriched in the modules M4_transcriptome_ and M10_transcriptome_ (**Figure 2C**). In addition, we observed that nearly all immune system related pathways were enriched in M3_transcriptome,_ the module which showed stepwise expression increase during ageing and lower transcript levels in G_DR nerves, reflected also in the enrichment of the pathways “Viral protein interaction with cytokine and cytokine receptor” and “cytokine-cytokine receptor interaction”, which contain a lot of cytokine, chemokines and their receptors (**Figure 2D**). Some specifically enriched pathways are worth noting, e.g. an enrichment of mTOR signalling in all upregulated proteins across all ageing cohorts, ECM-receptor interaction in M2_transcriptome_, FoxO and Apelin signalling in M2_proteome_ and AMPK signalling pathway, which was enriched in M6_proteome_, M10_transcriptome_ and the only the list of by DR upregulated transcripts/proteins (**Figure 2D**). Along the enrichment analyses for KEGG pathways and GO-Terms, we wanted to investigate which cell types may primarily contribute to the different co-expression modules in the transcriptome and proteome by performing an overrepresentation analysis (ORA) for all cell types of the intact sciatic nerve using cell-specific markers identified previously by single-cell RNAseq^30^. In both cases, the transcriptome and proteome exhibit similar cell type marker enrichment for some modules, yet also demonstrate different cell types to be overrepresented (**Figure 2E**): Significant ORA was identified for perineurial mesenchymal cells (pMES) in M1_transcriptome_; Schwann cells (SC) in M2_transcriptome_, M5_transcriptome_, M11_transcriptome_ and M5_proteome_; fibroblasts (Fb) and Mast cells (Mast) in M4_transcriptome_ and Fb only in M1_proteome_; endothelial cells (EC) as well as vascular smooth muscle cells and pericytes (vSMC_PC) in M9_transcriptome_ and M8_proteome_, reflecting their role in development and angiogenesis, GO-terms that were overrepresented in these modules (**Figure 2B**). Strikingly, the largest number of specific and mostly immune cell types were overrepresented in the M3_transcriptome_ (macrophages, mast cells, monocyte-derived dendritic cells, natural killer cells, proliferating mesenchymal cells, T cells) and M5_proteome_ (macrophages, Mast cells, monocyte-derived dendritic cells, natural killer cells and also Schwann cells) modules, indicating a substantial contribution of inflammation and immune cell infiltration to the induction of inflammatory processes during PNS ageing. However, Schwann cells were surprisingly also overrepresented in the “inflammation module” M5_proteome_, consistent with recent reports highlighting them as PNS-intrinsic immunocompetent glial cells^31,32^. Furthermore, the absence of overrepresentation of specific cell types in the “metabolism” modules (M6,M9_proteome_ and M10_transcriptome_, **Figure 2C**) indicates most likely an effect of DR on all cell types of the aged PNS (**Figure 2E**).

**Fig. 2.**
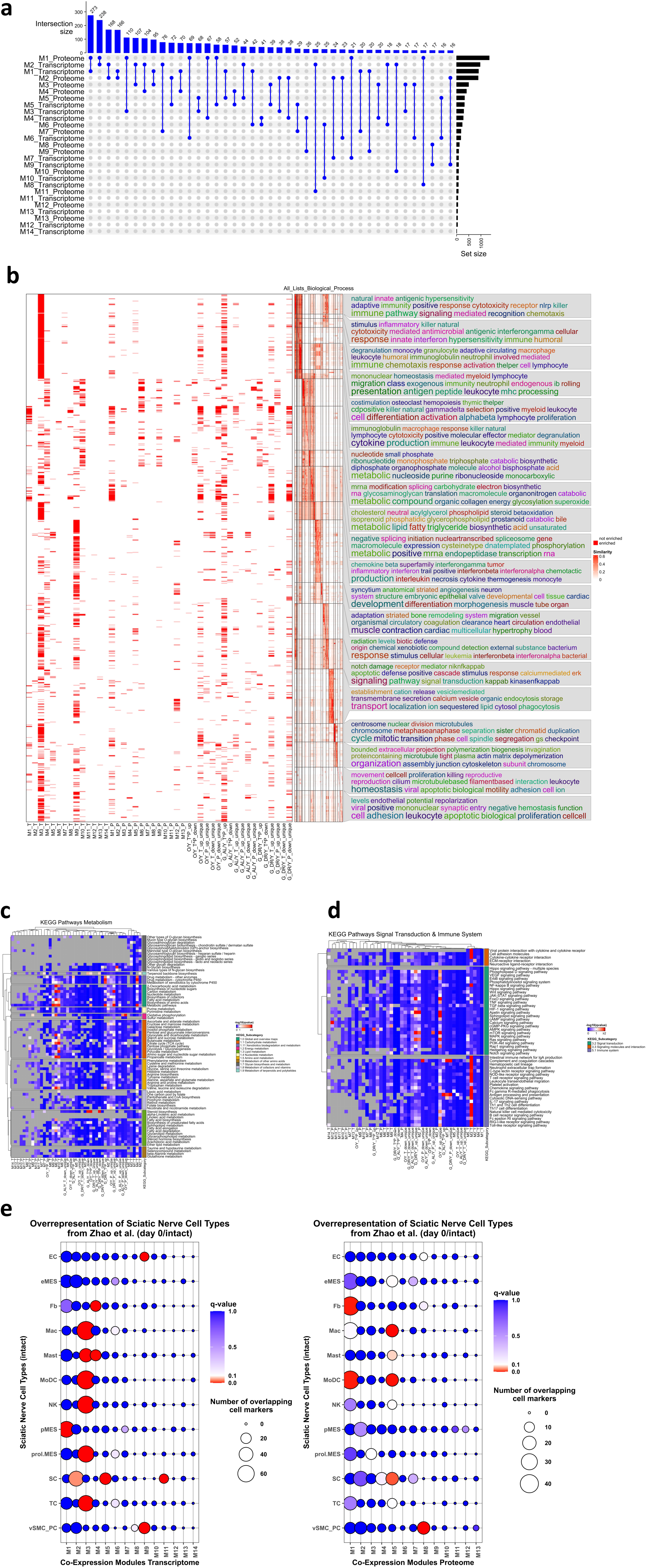
I Comparative and functional analysis of transcriptomic and proteomic changes of PNS ageing. **A** Upset plots showing the number of shared differentially regulated transcripts/proteins (depicted as intersection size) between different transcriptome and proteome co-expression clusters. The size of the respective module or gene/protein list is displayed on the right side of the plot (labelled as set list). **B** Gene ontology enrichment analysis of biological functions for all co-expression modules as well as all the different comparisons among all experimental groups. Semantically similar GO-terms are clustered together (the similarity matrix with specific clusters is depicted in the middle). A binary heatmap on the left shows if a GO-term was significantly enriched in the corresponding gene/protein list (columns). On the right word clouds are annotated to the semantic GO-term clusters with keywords that are overrepresented from all GO-terms located in this cluster, i.e. that serve as representative for the biological functions depicted in this cluster. **C, D** KEGG pathway enrichment heatmaps for all metabolism-related pathways (**C**) or immune system and signal transduction related pathways (**D**). In the columns are all gene/protein/module lists and in the rows are all pathways, split by KEGG subcategory. The q-values from the pathway enrichment are shown on a colour scale and gene/protein/module lists that show more similar pathway enrichment profiles cluster together. **E** Over-representation analysis of cell marker profiles in the intact sciatic nerve identified by scRNAseq^30^ across all co-expression modules of the transcriptome (left plot) and proteome (right plot). Abbreviations are similar to Zhao et al.^30^: EC: endothelial cells; eMES: endoneurial mesenchymal cells; Fb: Fibroblasts; Mac: Macrophages; Mast: Mast cells; MoDC: monocyte-derived dendritic cells; NK: natural killer cells; pMES: perineurial mesenchymal cells; prol.MES: proliferating mesenchymal cells; SC: Schwann cells; TC: T cells; vSMC_PC: vascular smooth muscle cells and pericytes

### Comparative and functional analysis of proteomic PNS injury profiles and their relationship to PNS ageing

After an injury, peripheral nerves undergo Wallerian degeneration, consisting of nerve fragmentation, inflammation and myelin clearance in the first 3 to 7 days after injury, after which regeneration is initiated in the subsequent weeks^33,34^. Previous work has indicated that low-grade inflammation reminiscent of that induced by injury is already present in intact old nerves and contributes to the age-related decline in peripheral nerve function^9^. Therefore, we investigated any potential similarities between processes occurring after injury and ageing-related changes. To this extent, we performed proteomics after peripheral nerve crush injury 3 days post crush injury (3dpc), 7dpc and 28dpc, to obtain molecular profiles during early and late nerve degeneration (3dpc and 7dpc) as well as early and advanced nerve regeneration (7dpc and 28dpc). Principal component analysis of the full datasets revealed a clear separation of crushed and intact nerves along the first principal component at all time points (**Extended Figure 4A, B**), with crushed nerves 28dpc being closest to intact nerves, reflecting the regenerative process (**Extended Figure 4B**). To obtain injury-specific proteome profiles at different time points, we identified subsets of proteins that were either shared-regulated, i.e. significantly up– or down-regulated at different time points post-injury, or uniquely regulated, i.e. up– or down-regulated at one time point but not the others. The largest subsets were the ‘pan-injury profiles’, i.e. those that were up– or down-regulated at all timepoints. This was followed by the subset of proteins that were up– or down-regulated at 7dpc and 28dpc, reflecting the induction of regenerative processes at 7dpc that extend to 28dpc and that are distinct to degenerative processes at 3dpc (**Extended Figure 4C**). A correlation network analysis of all ageing/DR and injury datasets revealed on the one hand a structured organization, i.e. post-injury datasets correlating positively with each other – with a stronger correlation between 7dpc and 28dpc as compared to 3dpc and 28dpc – and ageing datasets (old and geriatric groups) compared to young correlating positively with each other (**Figure 3A**). Notably, the G_AL/O dataset showed moderate positive correlations with all post-injury proteome datasets, suggesting that proteome changes at very old age partially converge with processes induced by PNS injury. By contrast, the G_DR/O dataset showed no correlation with post-injury proteomes, while the G_DR/G_AL dataset showed negative correlations with all post-injury proteome datasets. This indicates that the effects of dietary restriction are contrary to the processes occurring in mice at a very old (geriatric) age (**Figure 3A**). We then investigated the regulation of timepoint-specific and shared protein subsets across all ageing proteomes. Interestingly, higher protein levels were identified in G_AL and Y nerves for all pan-injury profiles, indicating that the injury-like proteome signature, which is induced at a very old (geriatric) age compared to an old age, is also active at young age during development and maturation (**Figure 3B**). Furthermore, dietary restriction prevented this induction and G_DR nerves exhibited the highest and lowest levels of proteins that were distinctly up– or down-regulated during late regeneration (28 dpc), suggesting that dietary restriction induces biological processes during late nerve regeneration (**Figure 3B**). To identify these processes, we conducted enrichment analyses (GO BP, GO CC and KEGG) for all post-injury proteome subsets. Several GO terms related to biological processes, such as ‘spliceosome’ and ‘translation’ (**Figure 3C**), as well as the GO_CC terms ‘spliceosomal’ and ‘ribosomal’, and ‘proteasome’ (**Extended Figure 4D**), were enriched in the list of proteins that were upregulated at all time points post-injury. Concomitant with this, the biological processes “cell cycle” and “mitosis” and the cellular component terms “chromosome proteins”, were also enriched in the list of upregulated proteins at all time points post-injury. This all together simply reflects the degradation and regenerative processes, including cell division, that occur after any tissue injury (**Figure 3C, Extended Figure 4D**). In addition, the enrichment of other structural, axonal, myelin and synaptic proteins (**Extended Figure 4D**), together with the biological processes of axonal transport, protein localisation and axon projection (**Figure 3C**), and the ECM-receptor interaction KEGG pathway (**Figure 3E**), in the downregulated proteins at all time points after injury can be considered a sign of general nerve degeneration and loss of tissue structure following nerve injury. In contrast, vesicle and lysosomal proteins, as well as ER and Golgi proteins, were enriched in the list of upregulated proteins (**Extended Figure 4D**). This is because, on the one hand, myelin degeneration and degradation occur after peripheral nerve injury, and, on the other hand, correct protein transport and modification are relevant for proper regeneration. Consequently, we also investigated whether signs of neurodegeneration and myelin degradation occur during ageing (**Extended Figure 5**): Indeed, we observed a loss of axonal and myelin proteins in geriatric nerves, while dietary restriction preserved axonal proteins and not only preserved, but also induced myelin proteins. This was accompanied by an increase in phagocytic and lysosomal proteins in geriatric nerves, whereas this increase could not be observed in G_DR nerves (**Extended Figure 5**). Another significant proportion of the enriched GO terms and KEGG pathways were related to metabolism. While most metabolic pathways, particularly those related to carbohydrate/sugar and fat metabolism, as well as the specific pathway “Valine, Leucine and Isoleucine Degradation”, were enriched in the list of downregulated proteins, primarily at the early time points 3dpc and 7dpc, they were only enriched in the list of upregulated proteins at 28dpc (**Figure 3C, 3D**). This occurred alongside the upregulation of mitochondrial and respiratory chain proteins at 28dpc only (**Extended Figure 4D** and **Supplementary Dataset 3**), which suggests that a metabolic switch occurs during late regeneration to help rebuild intact tissue integrity. Lastly, a major proportion of biological processes and pathways were related to immune system activation and inflammation. Both innate and adaptive immunity signalling pathways and biological functions/processes were represented (including viral defense responses (e.g. interferon-responsive/stimulated proteins), complement and antigen presentation and cGMP/PKG signalling). These were enriched in the list of all upregulated proteins (**Figure 3C, Extended Figure 4D** and **Supplementary Dataset 3**), reflecting the inflammation induced by injury, which is necessary for proper nerve degeneration and subsequent regeneration.

**Fig. 3.**
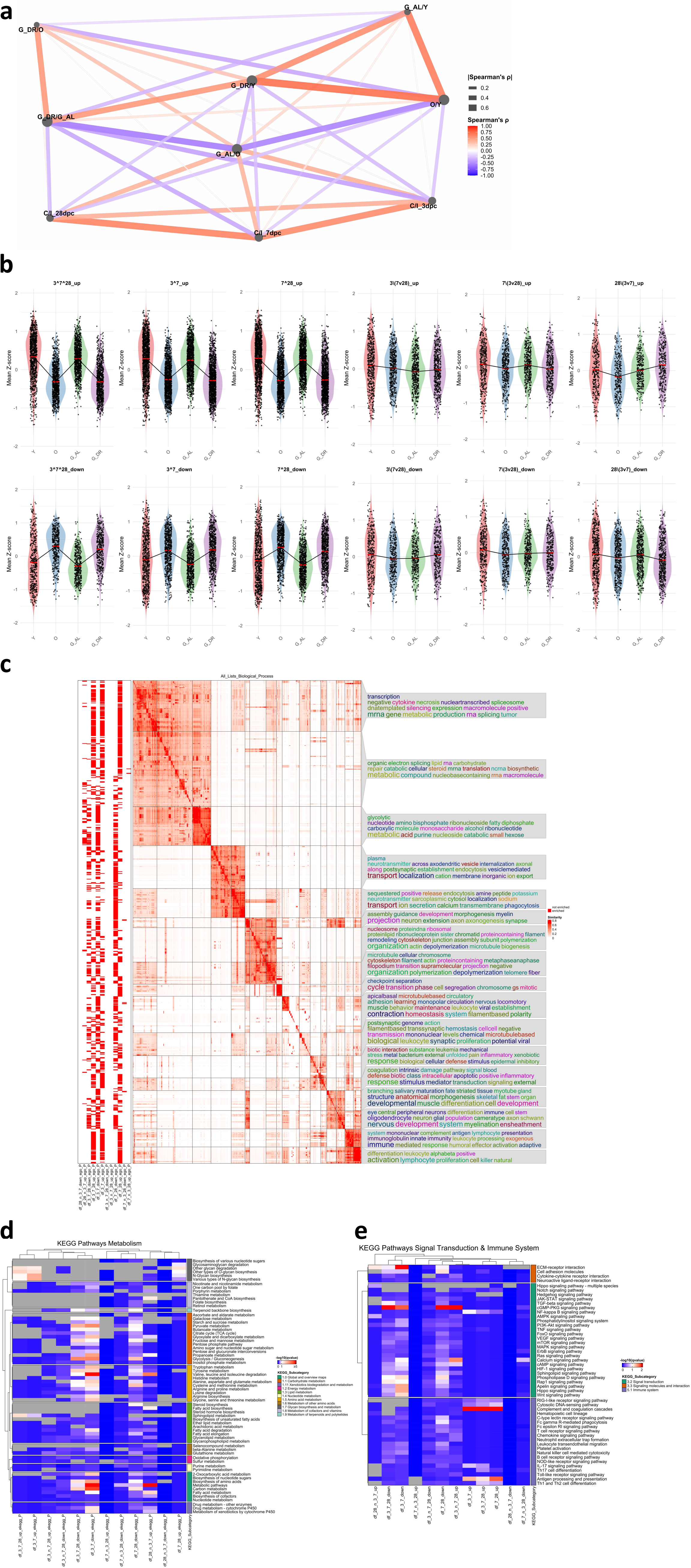
I Comparative and functional analysis of the proteomic response to PNS injury and ageing. **A** Correlation network of the Log_2_ fold changes of all full proteome datasets for all experimental comparisons of the ageing proteome, the 3dpc injury proteome and 7dpc/28dpc injury proteome. Spearman’s ρ is depicted on a colour scale as well as **I** Spearman’s ρ **I** is shown as a function of the line thickness. All correlation coefficients are provided in **supplementary table 2**. **B** Violin plots of the individual time-specific injury response proteome profiles during ageing and LT-DR. Shown are the mean-z-scores of all proteins of the respective injury response set for each experimental cohort (mean from n=6 mice per cohort). P-values represent FDR-adjusted p-values from pairwise Wilcoxon tests between the different cohorts. Logical operators “^” (and), “v” (or) and “/” (not in the following) are used to indicate the respective protein set as described in detail in the methods. **C** Gene ontology enrichment analysis of biological functions for all time-specific injury sets using semantic similarity analysis, methodically similar to Fig. 2. Column names for the different protein sets are described in detail in the methods. **D, E** KEGG pathway enrichment heatmaps for all metabolism-related pathways (**D**) or immune system and signal transduction related pathways (**E**) across all time-specific injury sets, methodically similar to Fig. 2.

### Identification of a type I interferon response signature during PNS ageing, dietary restriction and injury

In recent years, type I interferon responses (to either extracellular Interferon-α and –β or intracellular cGAS/STING signalling following the detection of dsDNA) have become recognised for their roles in mammalian ageing and senescence, particularly in the central nervous system (CNS)^35,36^. Additionally, well-documented side effects of interferon-α and –β treatment in humans include the development of peripheral neuropathy^37,38^. However, to our knowledge, changes in type I interferon signalling during peripheral nervous system (PNS) ageing or injury have not yet been described. Our previous enrichment analyses have associated immune and inflammatory transcript/protein changes in all datasets with the enrichment of type I interferon response pathways and predicted transcriptional regulators of the immune modules in the ageing/DR dataset were mostly found to functionally belong to the interferon response or regulatory system. Therefore, we specifically investigated the interferon system during peripheral nervous system (PNS) ageing and injury by mapping our datasets to previously described, evolutionarily conserved interferon network clusters^39^. Regulator and target proteins (i.e. interferon signature genes (ISGs)) from all five interferon network clusters were only upregulated in geriatric nerves, particularly in clusters 1 (C1) and 3 (C3) (**Figure 4A**). Interestingly, in geriatric nerves from caloric restriction mice, the ISG protein fold changes were reversed, i.e. downregulated, in the comparison to G_AL nerves but also to young nerves, indicating that dietary restriction prevents interferon network induction during PNS ageing, in particular for C3 (**Figure 4A**). One out of three key regulators of C3 is Stat1 (the other two, Usp18 and Irf9, were not detected in all proteomics datasets)^39^, which we identified to be upregulated in a stepwise manner during ageing and with complete prevention of induction by dietary restriction (**Figure 4A**). Strikingly, an interferon network signature with predominant C3 induction was also identified at 3dpc and 7dpc during the de– and early regeneration phases after peripheral nerve injury. However, at 28dpc, during the late regeneration phase, interferon network protein levels decreased markedly again. Concomitantly, Stat1 protein levels were significantly increased at 3dpc and 7dpc but returned to normal levels by 28dpc (**Figure 4A**). On the one hand, we wanted to validate the post-injury increase in Stat1 expression in an external dataset. On the other hand, we wanted to investigate Stat1 expression in individual cell types in the intact mouse sciatic nerve, as well as in the early phase after peripheral nerve injury. To this end, we re-analysed the scRNA-seq dataset by Zhao et al.^30^: In intact nerves, Stat1 is expressed by all cell types; however, it is primarily expressed by Schwann cells, endothelial cells, and macrophages (**Figure 4B**). Following injury, Stat1 is highly upregulated in immune cells, including macrophages, lymphocytes and granulocytes, at 1dpc, with continuous increased expression in these cell types, as well as in Schwann cells at 3dpc and 7dpc. This indicates that interferon-response signalling is induced in both immune cells and Schwann cells after peripheral nerve injury, as well as during ageing (**Figure 4B**).

**Fig. 4.**
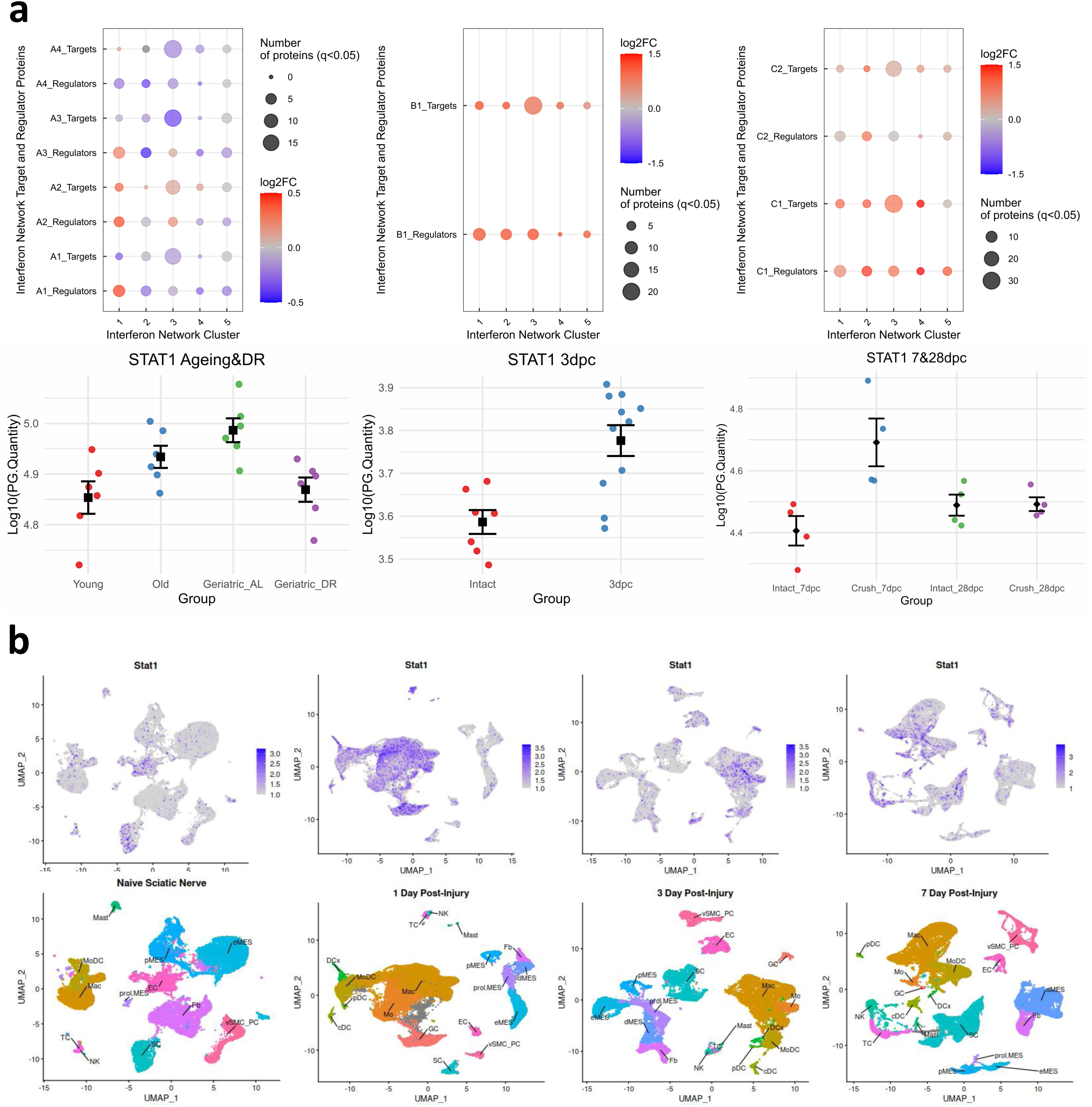
I Induction of Stat1 and an interferon response is a common denominator of age-and injury-associated peripheral nerve inflammation. **A** Mapping of significantly changed proteins to the IFN-network of regulators and interferon target genes as determined by Mostafavi et al.^39^. Left plot ageing&DR proteome for experimental comparisons A1 to A4 (A1 = O/Y, A2 = G_AL/Y, A3 = G_DR/Y, A4 = G_DR/G_AL), middle plot 3dpc proteome with B1 = 3dpc/Int, right plot 7&28dpc proteome with comparison C1 = 7dpc/Int, C2 = 28dpc/Int. Below each bubble plot are scatter plots of Log_10_(PG.Intensities) of Stat1, the main cluster 3 IFN-regulator^39^, shown. Q-values for the individual comparisons: A1: q = 0.00235, A2: q = 1.93E-07, A3: q = 0.268, A4: q = 3.78E-13, B1: q = 8.65E-21, C1: q = 5.487E-26, C2: q = 0.0037, for fold changes see **supplementary dataset 1**). **B** Re-analysis of Stat1 expression in a single-cell RNAseq dataset from^72^ during the early injury response. In the top plots are Stat1 levels after injury across all PNS cell types depicted, below are the annotations to the different cell types.

## Discussion

In this study, we performed transcriptomic and proteomic analyses of the sciatic nerves of young (3-month-old) and old (20-month-old) mice, as well as geriatric (24-month-old) mice that show advanced signs of PNS ageing and degeneration, as demonstrated in previous studies^7,9–12^. Furthermore, we analysed the sciatic nerves of mice that underwent lifelong caloric restriction, a known neuroprotective intervention, and identified major changes compared to the sciatic nerves of geriatric mice of the same age. Additionally, we obtained time-specific proteome profiles of sciatic nerve degeneration and regeneration following nerve crush injury. We then integrated these profiles with ageing and DR datasets, identifying similarities between ageing– and injury-induced biological processes and pathways.

Peripheral nerves exhibit impressive cellular heterogeneity, and balanced intercellular communication between Schwann cells, neurons, immune cells, and other stromal cells is necessary for maintaining peripheral nerves and their regenerative capacity. With advanced age, changes in and between all peripheral nervous system (PNS) cell types, especially Schwann cells^40^ and immune cells^7,9^, lead to nerve dysfunction, impaired regenerative capacity and the onset of age-associated peripheral neuropathies. Advanced age-related nerve pathology is present from around 24 months onwards in mice^7,10–12^. 18–20 months in mice corresponds to around 55–60 human years (HY) and 24+ months to 70+ HY^41^, which is exactly the age at which age-associated peripheral neuropathies begin to increase in prevalence (around 50–60 HY) and show a rapid increase from 70 HY onwards^1,42,43^. This indicates that major molecular changes occur during this period that ultimately lead to age-induced PNS dysfunction. Accordingly, we found that age-related molecular shifts occur in a stepwise fashion, with several linear and non-linear trajectories observed both at transcript and protein levels of old and geriatric nerves (**Figure 1**). This validates the previous observations of functional and cellular changes occurring during this period at a molecular level, and allows us to conclude that our datasets can be reliably used to investigate features of peripheral nervous system (PNS) ageing. Furthermore, in accordance with previous functional observations that dietary restriction can reverse or prevent age-associated PNS changes, we could also demonstrate that lifelong dietary restriction (DR) could ‘slow down’ PNS ageing. DR nerves were found to cluster together with old nerves in PCA (**Figure 1B, C**), and the fold changes in the DR dataset were found to be more strongly correlated with the fold changes in the old dataset than with the G_AL dataset from the same age group (ρ = 0.71 versus 0.56, respectively; **Figure 3A** and **Supplementary Table 2**). Moreover, we identified the induction of a validated senescence signature (SenMayo gene set^25^) as one typical hallmark of ageing^44^ in old nerves, with slightly stronger enrichment in geriatric nerves and similar enrichment in G_DR nerves (**Figure 1D**). These results also suggest that our datasets accurately reflect the molecular changes that occur in the ageing nerve in adaptation to DR, as demonstrated in previous functional and immunohistochemical studies^11,23^.

Peripheral nerve ageing is characterised by features similar to an injury response, including myelin degeneration and degradation^5,7^, inflammation^7,9^, and ECM remodelling^45^. Interestingly, we identified a positive correlation between ageing-induced and injury-induced changes in old and geriatric nerves, which was reversed in DR nerves (**Figure 3A** and **Supplementary Table 2**). Similar to the natural early-time course after injury, where myelin and injured axons degrade^45^ and phagolysosomal proteins are upregulated^5^, we identified that myelin and axonal proteins were downregulated and phagolysosomal proteins were upregulated only in the nerves of geriatric mice, whereas this did not occur in the nerves of geriatric mice with lifelong DR (**Extended Figures 3D and 5**). Metabolic dysregulation, characterised by mitochondrial dysfunction, decreased glucose tolerance and impaired fat oxidation, is widely regarded as a hallmark of ageing^44^. We identified several modules at the transcriptome (M10) and proteome (M6 and M9) levels that became upregulated in geriatric nerves and were further induced by dietary restriction (**Figures 1E** and **1F**). These modules showed strong enrichment in metabolic processes and pathways related to glucose and lipid catabolism, as well as oxidative phosphorylation and AMPK signalling was in addition enriched in the list of transcripts and proteins upregulated by DR (**Figures 2B, C, D**). Interestingly, the same metabolic processes and pathways were enriched in the lists of downregulated proteins at 3dpc and 7dpc, but became enriched again in the list of upregulated proteins at 28dpc, i.e. during late nerve regeneration (**Figure 3C, D**). Dietary restriction is known to improve glucose tolerance and utilisation at old age by stimulating AMPK signalling^46^, which shifts metabolism towards increased glucose catabolism and oxidative phosphorylation^46^. Furthermore, DR can regulate lipid catabolism by stimulating β-oxidation, which is believed to protect against age-related dysregulation of lipid metabolism^47^. In addition, restoration of fatty acid metabolism is a common denominator of geroprotective interventions, including dietary restriction, as demonstrated by a re-analysis of published ageing datasets in other organs than the PNS^48^. Following PNS injury, Schwann cells initially display a transient upregulation of glycolysis (during the first two days post-injury) to support the survival of injured axons. However, by 3 dpc (the time at which we obtained the proteomic profiles), glucose levels had returned to those observed in intact nerves^49^. Another study showed that at 7dpc, nerves displayed reduced oxidative phosphorylation and ATP-linked respiration, which began to recover at 28dpc^50^. Another landmark study by Sundaram et al. demonstrated that oxidative phosphorylation is induced over the course of nerve degeneration and regeneration, with the induction of mitochondrial respiration occurring between 14dpc and 28dpc and lipid oxidation potentially fuelling nerve regeneration and remyelination^51^. Therefore, it is tempting to hypothesise that the enrichment of glucose catabolism, oxidative phosphorylation, mitochondrial proteins, and lipid metabolism in the lists of proteins that are upregulated at 28dpc, as well as in the modules of DR-induced proteins (**Figures 1E, F, 2B, C, 3C, D**), is a sign of metabolic adaptation during ageing and DR, similar to the metabolic adaptation during late regeneration, which could explain partially the neuroprotective effects of DR seen in previous studies^11,23^. In addition, we found the degradation of branched-chain amino acids (BCAA), i.e. valine, leucine and isoleucine, to be enriched in the same modules and protein lists during ageing and after injury (**Figures 1E, F, 2B, C, 3C, D**). Given that BCAA restriction can increase healthspan and lifespan^52,53^ and contribute to healthy metabolism in old age, increased BCAA catabolism may be beneficial during ageing. While, to our knowledge, no studies have investigated the effects of BCAAs on peripheral nerve regeneration (PNS), a recent study demonstrated that restricting BCAAs led to reduced pain and inflammation, as well as increased nerve fibre density recovery after a plantar injury in mice^54^. This indicates potentially beneficial effects on nerve regeneration and regrowth that could contribute to the neuroprotective effects of DR.

Chronic inflammation during ageing, also known as “inflammageing”, is considered to be another hallmark of ageing^44^ and can also be observed during peripheral nerve ageing and after peripheral nerve injury^9^. In both datasets (the transcriptome and the proteome), we identified modules (M3_transcriptome_ and M5_proteome_) that became upregulated stepwise during ageing, with induction being prevented in DR nerves that showed strong enrichment in immune– and inflammation-related biological processes and signalling pathways (**Figures 2B, D**). Furthermore, immune cells such as macrophages and lymphocytes, as well as Schwann cells, were overrepresented in the above-mentioned modules. This indicates that various cell types contribute to PNS inflammation during ageing (**Figure 2E**). Concomitantly, immune and inflammatory pathways were also enriched in the lists of upregulated proteins at all timepoints after injury (**Figure 3C, E**). Specifically, NF-κB signalling and interferon response signalling were enriched in the previously mentioned modules. Transcription factor motif enrichment analysis also indicated an overrepresentation of type I interferon response signalling in the ‘inflammation modules’ induced during PNS ageing (**Figure 2B, Extended Figure 3A-C, Supplementary Table 1**). Recent work by Rasa et al. could demonstrate that *StatI*-mediated interferon-response signalling is a central characteristic of inflammageing and is counteracted by lifelong DR, which supports our results^55^. However, to our knowledge, no role for type I interferon response signalling, or indeed any role, has yet been described in the ageing PNS. We identified *Stat1* and an interferon network regulated by *Stat1*, which are induced in a stepwise manner during peripheral nerve ageing. Long-term dietary restriction prevents this induction. *Stat1* and the aforementioned interferon network are also induced during the degeneration and early regeneration phases (3dpc & 7dpc) after peripheral nerve injury. However, during the late regeneration phase (28dpc), protein levels and network activity return to normal (**Figure 4**), suggesting that *Stat1*-mediated interferon response signalling plays a role in both PNS ageing and degeneration/regeneration. Strikingly, recent work by Morozzi et al. has demonstrated that cGAS/STING is upregulated following peripheral nerve injury (PNS), triggering a type I interferon immune response^56^. Furthermore, Wang et al. have shown that cGAS upregulation in response to nerve injury is dependent on *StatI*^57^. Furthermore, perineural invasion occurs in multiple cancer types and Baruch et al. revealed a mechanism by which nerve injury leads to neuron-initiated, type I interferon-mediated inflammation to promote nerve regeneration^58^. While this inflammation is beneficial in the short term, it can contribute to the skewing of the tumour microenvironment and PD-1 resistance when it becomes chronic and PD-1 resistance can be reversed by knocking out the interferon-α receptor^58^. Zanon et al. demonstrated the beneficial role of type I interferon interferon-β, showing that treatment with interferon-β after nerve crush injury could promote nerve regeneration and motor recovery^59^. Furthermore, Barragan-Iglesias et al. demonstrated that type I interferons can act directly on nociceptors to produce pain sensitisation, which may be beneficial after injury to support healing^60^. However, long-term treatment with type I interferons, such as interferon-α for hepatitis C and interferon-β for multiple sclerosis, can lead to immune-mediated polyneuropathy^37,38,61^. Elevation of the chronic type I interferon response signalling is considered a key feature of inflammageing^44^, which can contribute to metabolic dysfunction^36^, neurodegeneration^62^ and cognitive decline^35^. Conversely, blocking chronic type I interferon signalling can reverse age-related cognitive dysfunction, as well as inflammation in the aged central nervous system (CNS)^35^, aged lungs^63^ and moreover, specifically knocking out *StatI* could prevent cellular ageing trajectories in the old brain^64^. Therefore, we can speculate that, while the type I interferon response after injury may promote nerve recovery, chronic induction, such as in aged nerves, may contribute to age-associated neuropathy and impaired nerve regeneration in old age. DR provides its neuroprotective effects, in part, by preventing the induction of the type I interferon response in the aged peripheral nervous system (PNS), as it does in other organs^55^.

These thoughts and our observations lead us to conclude that an old nerve is an injured nerve. This follows the recent thoughts of Mikolaj Ogrodnik that ageing in general can be considered a wound that never starts to heal^65^. This is analogous to the concept that tumours are wounds that attempted to start healing but did not heal^66^. This concept has also previously been suggested for schwannomas^67^, as these tumours often occur in areas of the body prone to lesions (e.g. the internal acoustic meatus) and also develop following peripheral nerve injuries in mice lacking the *Nf2* gene^68^. Moreover, recent work by Barrett et al. uncovered a nerve-injury-like state in human vestibular schwannomas with injury-like repair, Schwann cells and inflammation, which was associated with tumour formation and size^69^. Ogrodnik highlights that ageing induces a phenotype similar to that seen after an acute injury across basically all organs, and that aged tissue is ‘stuck’ in the ‘inflammation phase’ after injury, whereas tumours progress to the ‘proliferation phase’^65^. In addition, Ogrodnik suggests that the same processes and pathways that are detrimental during ageing are beneficial after injury and during development: This concept was introduced by George Williams in 1957 and is known as ‘antagonistic pleiotropy’ (AP)^70^. Indeed, our datasets reveal similar processes and pathways to be induced after injury and during ageing, including AP-1-associated transcription factor binding motifs (**Extended Figure 3C**, **Supplementary Table 1),** which have been previously linked to development and ageing across a large number of other murine tissues and cell types^71^. Furthermore, it is noteworthy that the injury-induced proteins that are downregulated in mature/old nerves and upregulated in geriatric nerves are also present at high levels in young nerves, providing further support for the AP hypothesis also in the context of PNS ageing.

Overall, we believe that our datasets support the idea that old nerves should be viewed as injured nerves. We identified several hallmarks of ageing^44^, such as inflammageing, metabolic dysregulation and induction of senescence, to be present in the old PNS. Ageing appears to induce a molecular phenotype in the PNS that is partially similar to that induced by injury, particularly with regard to metabolic and inflammatory changes. Furthermore, a well-established anti-ageing neuroprotective intervention — long-term dietary restriction — was found to partially reverse this phenotype. In summary, this article presents the most comprehensive molecular resource on peripheral nerve ageing, injury, and dietary modulation to date. We believe these datasets will serve as a valuable framework for understanding the molecular physiology of the ageing peripheral nervous system.

## Methods

### Experimental animals

All animals were on a C57BL/6 J background. Mice were group housed, with free access to standard chow and water and maintained with 12 h light and dark cycles. Temperature was 20 ± 2 °C during the experimental period. The animal procedures were performed according to the guidelines from Directive 2010/63/EU of the European Parliament on the protection of animals used for scientific purposes. Experiments were approved by the Thuringian State Office for Food Safety and Consumer Protection (licenses FLI-17-006, FLI-17-007 and FLI-18-017). Mice were considered to be geriatric at 24 months, old at 20 months and considered young at around 2-3 months. Dietary restriction was performed on female mice as previously described – sciatic nerves were taken from the same mouse cohort described by Rasa et al.^55^ – in brief, the daily food intake per animal was calculated on a daily basis and was the amount that refers to 70% for every animal dependent on its body weight for the G_DR cohort, G_DR and G_AL mice were single housed. Food was given once a day to the G_DR animals, starting from the age of 4 months, until 24 months, while G_AL animals had unlimited access (AL = *ad libitum*) to food during the experiment.

### Sciatic nerve crush injury

Unilateral injuries of sciatic nerves were performed with minimal invasion, as described previously ^9,68^. Briefly, mice were anaesthetized with isoflurane in oxygen, fur was removed with an electric razor (Aesculap ISIS, B. Braun AG, Melsungen Germany) and skin incised minimally. The biceps femoris muscle was separated to reveal the sciatic nerve. Using a smooth hemostatic forceps (width 0,6 mm, curved, clamping length 19 mm, Fine Science Tools GmbH, Heidelberg, Germany) the nerve was crushed mid-thigh. Muscle tissue was sutured using non-absorbable surgical suture material (Catgut GmbH, Markneukirchen, Germany) and skin closed with an AutoClip System (FST, 9 mm clips).

### RNA and protein isolation from sciatic nerve tissue

Frozen sciatic nerves, 1.4 mm ceramic (zirconium oxide) beads and peqGOLD TriFast™ were placed in a 2 ml tube and homogenized using a PeqLab-Precellys® 24 homogenizer. Samples were transferred to RNAse-free vessels, mixed with 0.25 ml chloroform for 10-15 sec and further incubated for 5 min at ambient room temperature. Centrifugation for 15 min at 12,000 x g at 4 °C produced a separation of two phases; the upper phase containing RNA and the lower phase containing the protein fraction. Proteins were obtained from the protein fraction by adding two volumes of isopropanol. Mixed solution was incubated for 10 min at room temperature or on ice, followed by 10 min centrifugation at 12,000 x g at 4 °C. The protein pellet was washed twice with 95 % ethanol before transfer to the FLI Core Facility Proteomics for further experiments. The RNA-containing aqueous layer was isolated into a fresh tube and combined with an equivalent volume of 70% ethanol to facilitate binding. Subsequent purification employed the spin-column protocol of the PureLink RNA Micro Kit (Thermo Fisher Scientific), adhering to the provided guidelines, including on-column treatment with the kit’s DNase mixture to eliminate genomic DNA. RNA was eluted in 18 μl distilled water and quality-checked initially using a NanoDrop spectrophotometer. Samples were then analyzed at the FLI Core Facility DNA Sequencing with Agilent’s 2100 Bioanalyzer to determine RNA integrity via the RNA integrity number (RIN) before RNA sequencing.

### RNA sequencing

Sequencing of RNA samples was performed using Illumina’s next-generation sequencing methodology^73^. In detail, total RNA was quantified and quality checked using Agilent 2100 Bioanalyzer Instrument (RNA 6000 Nano assay). Libraries were prepared using the TruSeq stranded mRNA library preparation kit (Illumina) according to the manufacturer’s instructions. Quantification and quality check of libraries was done using Agilent 2100 Bioanalyzer Instrument (DNA 7500 assay). Libraries were pooled and sequenced using three lanes of a HiSeq 2500 system running in 51 cycle/single-end/high-throughput mode. Sequence information was converted to FASTQ format using bcl2fastq v1.8.4. For expression analysis, the raw reads were quality-trimmed and filtered for low complexity with the tool preprocess from the SGA assembler (version 0.10.13, parameters –q 30 –m 50 –-dust)^74^. Passed reads were mapped to the mouse reference genome (GRCm38) with the Ensembl gene annotation (release 85) with TopHat2 (version 2.1.0)^75^ using conservative settings (--b2-sensitive –-no-coverage-search –-no-novel-juncs –-no-novel-indels –-transcriptome-max-hits=1). All reads mapping uniquely to an Ensembl gene were counted using FeatureCounts (multi-mapping or multi-overlapping reads were not counted, stranded mode was set to “–s 1”)^76^. The RNA sequencing data discussed in this publication have been deposited in NCBI’s Gene Expression Omnibus^77^ and are accessible through GEO Series accession number GSE137504 (young and old nerves) and GSE318229 (geriatric *ad libitum* and dietary restriction nerves).

### Proteomics of sciatic nerves – Ageing & DR experiments

#### Sample preparation and LC-MS/MS data independent acquisition/analysis (DIA)

The precipitated protein pellets were resuspended in lysis buffer (final concentration: 0.1 M HEPES/pH 8; 2% SDS; 0.1 M DTT), vortexed, then sonicated using a Bioruptor (Diagenode) (10 cycles of 1 min on and 30s off with high intensity @ 20°C). For reduction and full denaturation of the proteins, the lysates were first incubated at 95°C for 10 min and subsequently sonicated in the Bioruptor for a further 10 cycles, as before. The supernatants (achieved after centrifugation, 14,000 rpm, room temperature, 2 min) were then treated with iodacetamide (room temperature, in the dark, 30 min, 15 mM). After running a coomassie gel to estimate the amount of protein in each sample, approximately 60–120 μg of each sample were treated with 4 volumes ice-cold acetone to 1 volume sample and left overnight at −20°C, to precipitate the proteins. The samples were then centrifuged at 14,000 rpm for 30 min, 4°C. After removal of the supernatant, the precipitates were washed twice with 400 μL 80% acetone (ice-cold). Following each wash step, the samples were vortexed, then centrifuged again for 2 min at 4°C. The pellets were allowed to air-dry before dissolution in digestion buffer (60 μL, 3 M urea in 0.1 M HEPES, pH 8) with sonication (3 cycles in the Bioruptor as above) and incubation for 4 h with LysC (1:100 enzyme: protein ratio) at 37°C, with shaking at 600 rpm. The samples were then diluted 1:1 with milliQ water (to reach 1.5 M urea) and incubated with trypsin (1:100 enzyme: protein ratio) for 16 h at 37°C. The digests were acidified with 10% trifluoroacetic acid, then desalted with Waters Oasis® HLB μElution Plate 30 μm in the presence of a slow vacuum. This process involved conditioning the columns with 3 × 100 μL solvent B (80% acetonitrile; 0.05% formic acid) and equilibration with 3 × 100 μL solvent A (0.05% formic acid in milliQ water). The samples were loaded, washed 3 times with 100 μL solvent A, then eluted into PCR tubes with 50 μL solvent B. The eluates were dried down with the speed vacuum centrifuge and dissolved in 50 μL 5% acetonitrile, 95% milliQ water, with 0.1% formic acid prior to analysis by LC–MS/MS. Approximatively 1μg of reconstituted were separated using a nanoAcquity UPLC (Waters, Milford, MA) coupled to an Orbitrap Fusion Lumos (Thermo Fisher Scientific, Bremen, Germany). Peptide mixtures were separated in trap/elute mode, using a trapping (Waters nanoEase M/Z Symmetry C18, 5μm, 180 μm x 20 mm) and an analytical column (Waters nanoEase M/Z Peptide C18, 1.7μm, 75μm x 250mm).. The outlet of the analytical column was coupled directly to an Orbitrap Fusion Lumos mass spectrometers (Thermo Fisher Scientific, San Jose, CA) using the Proxeon nanospray source. Solvent A was water, 0.1% formic acid and solvent B was acetonitrile, 0.1% formic acid. The samples were loaded with a constant flow of solvent A, at 5 μL/min onto the trapping column. Trapping time was 6 min. Peptides were eluted via the analytical column with a constant flow of 300 nL/min. During the elution step, the percentage of solvent B increased in a nonlinear fashion from 0% to 40% in 120 min. Total runtime was 145 min, including cleanup and column re-equilibration. The peptides were introduced into the mass spectrometer via a Pico-Tip Emitter 360 µm OD x 20 µm ID; 10 µm tip (New Objective) and a spray voltage of 2.2 kV was applied. The capillary temperature was set at 300 °C. The RF lens was set to 30%. Full scan MS spectra with mass range 350-1650 m/z were acquired in profile mode in the Orbitrap with resolution of 120,000 FWHM. The filling time was set at maximum of 20 ms with an AGC target of 5 × 10^5^ ions. DIA scans were acquired with 40 mass window segments of differing widths across the MS1 mass range. The HCD collision energy was set to 30%. MS/MS scan resolution in the Orbitrap was set to 30,000 FWHM with a fixed first mass of 200m/z after accumulation of 1 × 10^6^ ions or after filling time of 70ms (whichever occurred first). Data were acquired in profile mode. For data acquisition and processing Tune version 2.1 and Xcalibur 4.0 were employed.

### Proteomic data processing

DIA raw data were analyzed using the directDIA pipeline in Spectronaut v.17 (Biognosys, Switzerland) with BGS settings besides the following parameters: Protein LFQ method= QUANT 2.0, Proteotypicity Filter = Only protein group specific, Major Group Quantity = Median peptide quantity, Minor Group Quantity = Median precursor quantity, Data Filtering = Qvalue, Normalizing strategy = Local Normalization. The data were searched against a species specific (Mus musculus, 16,747 entries, v. 160106) and a contaminants (247 entries) Swissprot database. The identifications were filtered to satisfy FDR of 1 % on peptide and protein level. Relative protein quantification was performed in Spectronaut using a pairwise t-test performed at the precursor level followed by multiple testing correction according to Benjamini-Hochberg. The data (candidate table) and data reports (protein quantities) were then exported for further data analyses. The obtained dataset has been deposited on the MassIVE repository (MSV000100637).

### Proteomics of sciatic nerves – Injury experiments

#### Sample preparation

Samples were homogenized and lysis buffer (fc 4% SDS, 100 mM HEPES, pH 8.5, 50 mM DTT) was added to 15 µg of each sample. Samples were then boiled at 95°C for 10 min and sonicated using a tweeter. Reduction was followed by alkylation with 200 mM iodoacetamide (IAA, final concentration 15 mM) for 30 min at room temperature in the dark. Samples were acidified with phosphoric acid (final concentration 2.5%), and seven times the sample volume of S-trap binding buffer was added (100 mM TEAB, 90% methanol). Samples were bound on 96-well S-trap micro plate (Protifi) and washed three times with binding buffer. Trypsin in 50 mM TEAB pH 8.5 was added to the samples (1 µg per sample) and incubated for 1 h at 47°C. The samples were eluted in three steps with 50 mM TEAB pH 8.5, elution buffer 1 (0.2% formic acid in water) and elution buffer 2 (50% acetonitrile and 0.2% formic acid). The eluates were dried using a speed vacuum centrifuge (Eppendorf Concentrator Plus, Eppendorf AG, Germany) and stored at –20° C. Before analysis, samples were reconstituted in in MS Buffer (5% acetonitrile, 95% Milli-Q water, with 0.1% formic acid), spiked with iRT peptides (Biognosys, Switzerland) and loaded on Evotips (Evosep) according to the manufacturer’s instructions. In short, Evotips were first washed with Evosep buffer B (acetonitrile, 0.1% formic acid), conditioned with 100% isopropanol and equilibrated with Evosep buffer A. Afterwards, the samples were loaded on the Evotips and washed with Evosep buffer A. The loaded Evotips were topped up with buffer A and stored until the measurement.

### LC-MS/MS data independent acquisition/analysis (DIA)

Peptides were separated using the Evosep One system (Evosep, Odense, Denmark) equipped with a 15 cm x 150 μm i.d. packed with a 1.5 μm Reprosil-Pur C18 bead column (Evosep performance, EV-1137, Denmark). The samples were run with a pre-programmed proprietary Evosep gradient of 44 min (30 samples per day) using water and 0.1% formic acid and solvent B acetonitrile and 0.1% formic acid as solvents. The LC was coupled to an Orbitrap Exploris 480 (Thermo Fisher Scientific, Bremen, Germany) using PepSep Sprayers and a Proxeon nanospray source. The peptides were introduced into the mass spectrometer via a PepSep Emitter 360-μm outer diameter × 20-μm inner diameter, heated at 300°C, and a spray voltage of 2 kV was applied. The injection capillary temperature was set at 300°C. The radio frequency ion funnel was set to 30%. For DIA data acquisition, full scan mass spectrometry (MS) spectra with a mass range of 350–1650 m/z were acquired in profile mode in the Orbitrap with a resolution of 120,000 FWHM. The default charge state was set to 2+, and the filling time was set at a maximum of 20 ms with a limitation of 3 × 10^6^ ions. DIA scans were acquired with 40 mass window segments of differing widths across the MS1 mass range. Higher collisional dissociation fragmentation (normalized collision energy 30%) was applied, and MS/MS spectra were acquired with a resolution of 30,000 FWHM with a fixed first mass of 200 m/z after accumulation of 1 × 10^6^ ions or after filling time of 45 ms (whichever occurred first). Data were acquired in profile mode. For data acquisition and processing of the raw data, Xcalibur 4.4 (Thermo) and Tune version 4.0 were used.

### Proteomic data processing

DIA raw data were analyzed using the directDIA pipeline in Spectronaut v.20 (Biognosys, Switzerland) with BGS settings besides the following parameters: Protein LFQ method= QUANT 2.0, Proteotypicity Filter = Only protein group specific, Major Group Quantity = Median peptide quantity, Minor Group Quantity = Median precursor quantity, Data Filtering = Qvalue, Normalizing strategy = Local Normalization. The data were searched against a species specific (Mus musculus, 16,747 entries, v. 160106) and a contaminants (247 entries) Swissprot database. The identifications were filtered to satisfy FDR of 1 % on peptide and protein level. Relative protein quantification was performed in Spectronaut using a pairwise t-test performed at the precursor level followed by multiple testing correction according to Benjamini-Hochberg. The data (candidate table) and data reports (protein quantities) were then exported for further analyses. Datasets from 7dpc and 28dpc were analysed in a similar manner to the 3dpc dataset and were previously deposited on the PRIDE database (PXD041026)^79^ as well as partially described in a previous publication^45^. The obtained 3dpc dataset has been deposited on the MassIVE repository (MSV000100636).

### Bioinformatic data analysis

All data analysis steps were completed with R^78^. Data wrangling steps were performed using primarily dplyr functions from the tidyverse package^79^. Venn diagrams were made using the VennDiagram package^80^. PCA and further clustering analysis was performed on normalized counts after transformation using DESeq2 vst() function for transcriptome data and log_2_ transformed PG.Quants using the prcomp function and visualization was performed using either DESeq2 package^81^ for transcriptome data or the factoextra package for proteomics data. For differential expression analysis of transcriptome data, DESeq2 package was used. Rows/genes with only zero counts in all samples were filtered out before proceeding with the differential expression analysis. Furthermore, the “apeglm” R package^82^ was used for fold change shrinkage. Transcripts were considered differentially expressed with q ≤ 0.05. Differentially expressed proteins were obtained by using a q ≤ 0.05 cutoff based on Spectronaut analyses as described before. The following experimental comparisons of interest were defined for the ageing dataset: 1 = Old/Young, 2 = Geriatric_AL/Young, 3 = Geriatric_DR/Young, 4 = Geriatric_DR/Geriatric_AL, 5 = Geriatric_AL/Old, 6 = Geriatric_DR/Old and labelled as indicated in the respective figure legends. Weighted gene/protein correlation network analysis (WGCNA) was performed using the WGCNA^83^ and CEMiTool^84^ R packages. In brief, to reduce noise and increase cluster specificity, as suggested previously^84–86^, all in at least one of the aforementioned experimental comparisons significantly changed transcripts/proteins (q ≤ 0.05) were included in the WGCNA analysis. First, clustering of module eigengenes and visualization with a dendrogram was performed to identify a biologically reasonable threshold, for both the transcriptome and the proteome a threshold of 0.7 eigengene correlation was used. Subsequently, signed networks were constructed for better biological interpretability and biweight midcorrelation was used as a robust correlation measure. The final WGCNA function with previously mentioned settings was in the end run using the cemitool() function that chooses the soft-thresholding power β automatically to achieve scale-free topology^84^. Gene set enrichment analysis (GSEA) was performed with the “fgsea” package^87^ and were visualized using the gggsea package^88^. For senescent phenotype analysis on transcriptome level the “SenMayo” gene set for mice was used from Saul et al.^25^. Heatmaps were created using the “ComplexHeatmap” package^89^ of row-wise z-score normalized values from the respective experiment (log_2_ PG.Quants or vst-transformed counts). The resulting dendrogram was then resorted with “dendsort” package^90^. Mapping of the proteome to the transcriptome for comparative analysis was done with genenames using the genekitr package^91^ and UniProt.ws package. Analysis of module/gene product list similarity with UpSet plots was performed using the UpSetR package^92^ as indicated in the respective figures. Gene ontology and KEGG pathway enrichment analyses were conducted using the clusterProfiler R package^93^ based on an over-representation analysis (ORA). GO-network function visualization using an enrichment map was done using the clusterProfiler R package^93^. Visualization of significantly enriched GO-terms (q < 0.1) and clustering was done using the “simplifyEnrichment” package^94^ with kmeans clustering as method for semantic similarity clustering. Identification of significantly over-represented cell types in the WGCNA modules was done using the scRNAseq-dataset of intact sciatic nerves (day_0) by Zhao et al.^30^, which was retrieved from GEO database (GSE198582), pre-processed as stated in the publication by Zhao et al.^30^ and subsequently the top150 marker genes identified using the FindAllMarkers function from the Seurat package (V4)^95^. Over-representation analysis was done using the phyper function (phyper(q – 1, m, n, k, lower.tail = FALSE)) with q = as the overlap between the respective module and cell type, m = as all gene products of the respective module, n = all gene products minus m and k = the overlap between the respective cell type set and all gene products of the respective dataset, ORA results were plotted using ggplot2. Identification of module-specific transcriptional regulators by transcription factor motif enrichment analysis was done using the RcisTarget package as described previously^96^. The database used in this paper was the mm10_refseq_r80_v10 with ± 10kbp around the transcription start site from the cisTarget resources website^97^. To increase specificity, the cutoff (aucMaxRank) was set to 0.05. Top15 motifs for module M3_transcriptome_ and M5_proteome_ were visualized using ggplot2 as well as in **supplementary table 1** and results for all motifs are reported in **supplementary dataset 2**. Injury proteomics datasets were in general analysed as described above for the ageing proteomics data. Here the relevant experimental comparisons were for the 3dpc dataset: 3dpc_Crush/3dpc_Intact and for the 7/28dpc dataset: 7dpc_Crush/7dpc_Intact and 28dpc_Crush/28dpc_Intact. Correlation network using UniProt IDs was computed using the correlate()(function from the “corrr” package and the network was visualised by creating an igraph object^98^ and then using the ggraph package for visualization. Injury profiles were identified by data wrangling to get lists of shared and unique proteins across the different time points (3’7’28 up/down = sharedly up/downregulated across all time points; 3’7 up/down and 7’28 up/down = sharedy up/downregulated 3dpc and 7dpc or 7dpc and 28dpc, respectively; 3\\(7v28) up/down = uniquely up/downregulated at 3dpc and not at 7dpc or 28dpc, similar notation applies to 7\\(3v28) and 28\\(3v7)). Due to problems with plotting special characters, group set labels in heatmaps are modified, e.g. the set 3’7’28 up becomes “df_3_7_28_up” and the set 3\\(7v28) up becomes “df_3_n_7_28_up”. Interferon network analysis was performed as previously described^99^ using the network clusters identified by Mostafavi et al.^39^. UMAP plots of Stat1 during PNS degeneration were done using the RShiny database from the injured sciatic nerve atlas (iSNAT)^30^. The myelin protein set was retrieved from^45^ and proteins annotated to GO-terms axon, phagocytic vesicle and lysosome were retrieved using the biomaRt package^100^. All z-scored protein/transcript intensities as well as files with the results from the differential expression analysis for all datasets are provided in **supplementary dataset 1**.

### Use of artificial intelligence (LLMs) for manuscript preparation

During the preparation of this work, the authors used OpenEvidence, ChatGPT and DeepL for literature search (OE and CG) and to improve language and readability of the content (CG and DL). After using these tools, the authors reviewed and edited the content as needed and take full responsibility for the content of the publication.

### Statistical procedures

All analyses were performed using Graphpad Prism 8.4 or R^78^. Shapiro-Wilk normality test was carried out before further statistical analysis was performed. Depending on the result, parametric (RM-ANOVA with post-hoc paired t-tests) or non-parametric (Friedman-test with post-hoc Dunn’s multiple comparison test) statistical tests were conducted, as stated in the figure legends, p-values (p) were subsequently FDR-corrected. Statistical tests for proteomic analysis and functional annotation and enrichment analyses were performed as indicated in the respective material and methods section. All data are presented as mean ± SEM. Statistical significance was accepted if p ≤ 0.05, unless otherwise stated in the respective figure legend.

## Acknowledgments

The authors would like to thank Rose-Marie Zimmer and Norman Rahnis for excellent technical assistance. Furthermore, the authors would like to thank Madelaine Braune and Claudia Maisch for animal husbandry. The Core Facility DNA sequencing (Cornelia Luge, Marco Groth) of the FLI is gratefully acknowledged for their technological support in library preparation and sequencing. Open access funding provided by Projekt DEAL.

